# *SRF* fusion oncogenes encode constitutively activated chimeric transcription factors in myoid soft tissue tumors

**DOI:** 10.1101/2025.09.18.676997

**Authors:** Constance Pirson, Ariane Sablon, Axelle Loriot, Pierre Autin, A. Koen Braat, Marie Karanian, Cristina R Antonescu, Franck Tirode, Jean-Baptiste Demoulin

## Abstract

SRF fusion genes drive the pathogenesis of muscle-related soft tissue tumors, including subsets of perivascular tumors, inflammatory myofibroblastic tumor, and rhabdomyosarcoma. SRF encodes Serum Response Factor, a well-characterized transcription factor that regulates muscle development. We characterized four fusion genes: SRF::RELA, SRF::FOXO1, SRF::ICA1L, and SRF::PDGFRB. All localized to the nucleus and dimerized through the SRF MADS box. SRF::RELA, SRF::FOXO1, and SRF::ICA1L acted as constitutively active transcription factors independent of canonical cofactors, binding SRF target promoters and driving transcription via the partner transactivation domain (TAD). A cryptic TAD was uncovered in ICA1L. These fusions promoted mesenchymal cell growth and upregulated muscle-related genes in mesenchymal stem cells, recapitulating transcriptional signatures of human tumors. In contrast, SRF::PDGFRB acted through its kinase domain, was imatinib-sensitive, activated STAT1 and stimulated inflammation genes, consistent with its tumor phenotype. SRF fusions thus define a novel family of oncogenes in human myoid soft tissue tumors.

**Graphical Abstract:** 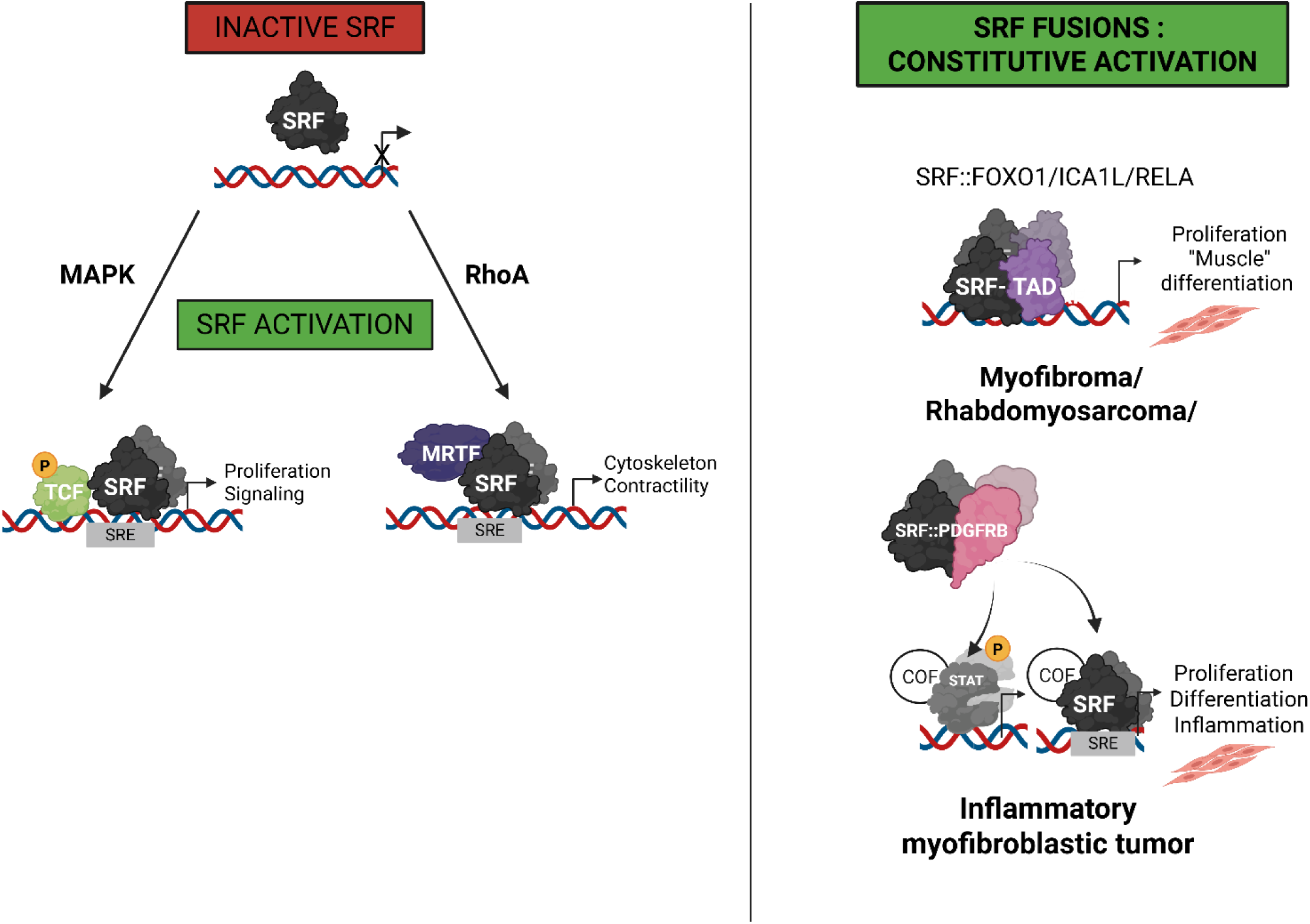

## INTRODUCTION

Serum Response Factor (SRF) is a key transcription factor that controls proliferation and differentiation of muscle cells. SRF regulates gene transcription through its DNA-binding domain, also named the MADS box, which binds to serum response elements (SRE) in target gene promoters [1]. Two distinct signaling pathways can activate SRF-mediated transcription, depending on the cofactors involved [2]. In the proliferation pathway, SRF interacts with ternary complex factors (TCF), such as ELK1, which are phosphorylated and activated by (MAPK) in response to growth factors. These cofactors drive the transcription of genes related to cell growth and the immediate early response. In the differentiation pathway, myocardin-related transcription factors (MRTF), such as MRTF-A and -B, are activated by cytoskeletal changes via RhoA GTPase signaling and actin polymerization. MRTFs bind to SRF and regulate genes involved in muscle cell differentiation. TCF and MRTF cofactors compete for binding to the SRF MADS box. By limiting MRTF access to SRF, TCFs suppress contractile gene expression and promote proliferation [2, 3].

Although many studies have linked SRF signaling to cancer promotion, cell invasion and angiogenesis [4–7], alterations of the *SRF* gene have only been identified recently [8]. We and others reported *SRF* rearrangements in 56 cases of myoid soft tissue tumors, including perivascular tumors, myofibroblastic tumors and rhabdomyosarcomas (Table 1) [9–17]. Most of these fusions were identified in a new group of benign perivascular tumors, characterized by myoid or immature smooth muscle differentiation, now recognized as a separate entity distinct from myofibroma and myopericytoma [18]. In addition, several *SRF* fusions were reported in a small subset of rhabdomyosarcoma, characterized by a favorable outcome compared to other rhabdomyosarcoma subtypes [19]. Finally, *SRF::PDGFRB* was described in one case of inflammatory myofibroblastic tumor (IMT), which is a rare, locally aggressive neoplasm, characterized by myofibroblastic differentiation accompanied by an abundant inflammatory infiltrate [20].

**Table 1.** SRF fusions in soft tissue tumors. Fusions reported in the literature (May 2025).

| 5' partner genes | 3' partner genes | Diagnosis | Cases | Ref. |
| --- | --- | --- | --- | --- |
| **Fusions with a transcription factor** |  |  |  |  |
| SRF exon4 | RELA exon11 | Cellular Myofibroma/myopericytoma | 8 | [10], [52] |
| SRF exon3/4/5 | RELA exon2/7/9 | Perivascular tumors | 9 | [16],[53],[54],[55] |
| SRF exon4/5 | RELA exon9 | Uterine tumor with myogenic differentiation | 3 | [56] |
| SRF exon3 +100pb<br>intron3 | E2F1 exon5 -66bp | Myoepithelioma | 2 | [11] |
| SRF exon6 | FOXO1 exon2 | Rhabdomyosarcoma | 1 | [13] |
| SRF exon5 | STAT6 exon17 | Myofibroma | 1 | [15] |
| SRF exon6 | MKL2 exon12 | Undifferentiated soft tissue sarcoma | 1 | [57] |
| **Fusions with a transcription cofactor** |  |  |  |  |
| SRF exon4 | NCOA1 exon13 | Rhabdomyosarcoma | 2 | [13] |
| SRF exon6 | NCOA2 exon11/12 | Spindle cell rhabdomyosarcoma | 2 | [9],[58] |
| SRF exon2/4 | NCOA2 exon12/14/15 | Perivascular tumors | 7 | [16],[54],[55] |
| SRF exon4 | NCOA3 exon12 | Cellular myofibroma | 1 | [53] |
| SRF exon2/4/5 | CITED1 exon2/3 | Perivascular tumors | 6 | [16],[17],[55] |
| **Fusions with a cytoplasmic protein** |  |  |  |  |
| SRF exon2/4/5 | ICA1L exon10/11/12/13 | Cellular myoid tumors | 8 | [12],[17],[55] |
| SRF ex4 | C3orf62 ex3 | Perivascular tumors | 2 | [55],[10] |
| SRF exon6 -41bp | PDGFRB exon12 -34bp | Inflammatory myofibroblastic tumor | 1 | [14] |
| SRF exon4 -3bp/6 | NFKBIE exon1 -235bp | Perivascular tumors | 2 | [16],[53] |

SRF fusion genes are in frame and code for chimeric proteins that have not been characterized yet. All fusions retain at least the two first exons of SRF, which encode the MADS box. Most 3’ fusion partners are transcriptional regulators (Table 1). Their transactivation domain (TAD) is always retained, creating putative chimeric transcription factors. However, some partners are devoid of known transcriptional function, such as ICA1L, C3orf62, NFKBIE and PDGFRβ, encoded by *PDGFRB*.

Comprehensive functional studies of these SRF fusion proteins are lacking. The goal of the present study was to elucidate the molecular mechanisms activating the fusions. We focused on four fusions, including SRF::RELA, SRF::FOXO1, SRF::ICA1L and SRF::PDGFRβ. We showed that SRF fusions unleash transcriptional activity, drive cell proliferation, and alter gene expression profiles, consistent with their oncogenic potential.

## RESULTS

### SRF fusions exhibit constitutive transcriptional activity

To understand the molecular and cellular mechanisms of transformation, we focused on the selected four fusion variants (Fig. 1A). *RELA* is the most recurrent partner, with fusions typically including the first four exons of *SRF* with varying numbers of *RELA* exons. We studied a short (S) and a long (L) version of the SRF::RELA fusion, which encompasses exon 11 only or exons 2 to 11 of *RELA*, respectively. Only the long version includes the Rel homology domain (RHD) of *RELA*, required for DNA binding and NF-κB transcriptional activity, whereas the short version lacks this domain. Additionally, we selected the SRF::FOXO1 fusion, described in rhabdomyosarcoma, and two fusions with non-transcriptional partners: SRF::ICA1L, the second most recurrent fusion, and SRF::PDGFRβ.

**Figure 1.**
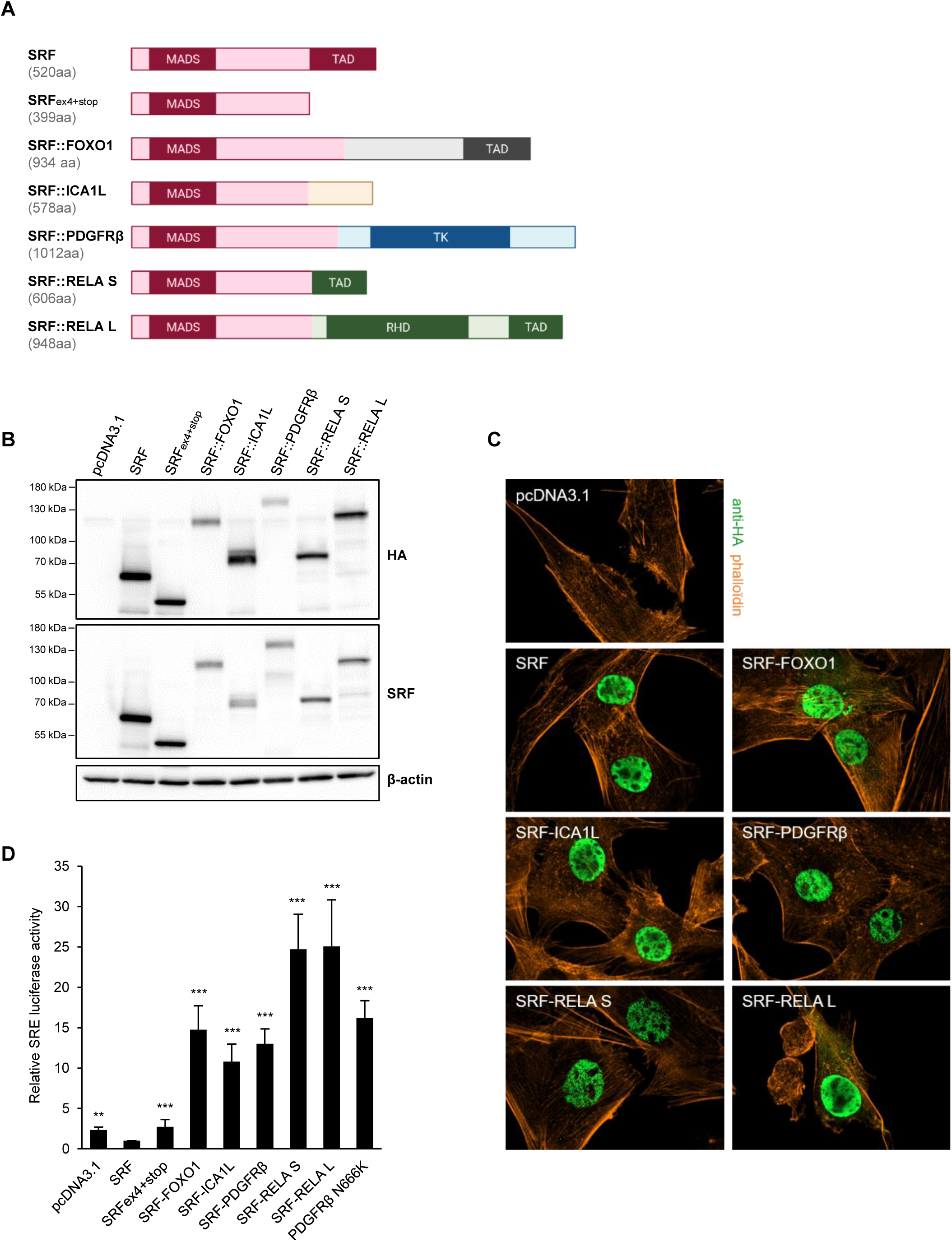
Expression, localization and transcriptional activity of SRF fusions. (A) Schematic structures of the studied SRF fusion proteins. SRFex4+stop was added as a control. (B) C2C12 cells were transfected with the indicated SRF fusions, the expression of which was analyzed by western blot with anti-HA and anti-SRF antibodies (N=5). (C) Subcellular localization of SRF fusions was analyzed by immunofluorescence with anti-HA antibody (N=3). (D) C2C12 were co-transfected with SRF fusions, SRE-luciferase reporter vector and an internal control. After 24h of transfection, luminescence was measured in cell lysates. The mean of three independent experiments is represented with SEM (One-way ANOVA test with *p < 0.05; **p < 0.01; ***p < 0.001 compared to SRF WT). The empty vector pcDNA3.1 was used as negative control.

First, we determined the expression levels of SRF fusions by western blotting in the transiently transfected C2C12 myoblast cell line. All fusions were correctly expressed at the expected size, though at a slightly weaker level than SRF WT (Fig. 1B). Immunofluorescence showed that, like SRF WT, all fusions localized predominantly in the nucleus (Fig. 1C). Accordingly, fusion proteins were present in nuclear extracts of transfected cells, but not in cytosolic fractions, as shown by western blot after cell fractionation (Supplementary Fig. S1).

Next, transcriptional activity of SRF fusions was measured using a SRE luciferase reporter assay. SRF fusions significantly activated SRE reporter by at least 10-fold, whereas SRF WT consistently decreased luciferase activity (Fig. 1D). PDGFRβ N666K, an oncogenic mutant reported in myofibroma and known to activate SRE transcriptional activity, was used as positive control [25]. Finally, a construct containing only the first four exons of SRF (SRF ex4+stop) lacked transcriptional activity, underlying the essential role played by the partner gene. In parallel, NF-κB transcriptional activity was also assessed using a dedicated reporter assay. SRF::RELA^L^ significantly activated the NF-κB reporter, consistent with the presence of the RHD domain, whereas SRF::RELA^S^ showed no activity (Supplementary Fig. S2). In conclusion, our data demonstrated that SRF fusion gene products are nuclear proteins with constitutively activated transcriptional activity.

### SRF fusions rely on DNA binding, except for SRF::PDGFRβ

We next investigated the role of the MADS box, common to all SRF fusions (Fig. 2A). We first introduced the S162D mutation, which is known to disrupt DNA binding by adding a negative charge that repels DNA [21]. Except for SRF::PDGFRβ, S162D mutants failed to induce SRE reporter activity, highlighting the key role of the MADS box/SRE interaction (Fig. 2B). We also introduced the V194E substitution to assess the role of SRF cofactors. Indeed, Val194, located in the second β-strand of the MADS box, is critical for binding TCF and MRTF cofactors, including Elk-1 and myocardin [2]. However, SRE luciferase assay suggested that the Val194 residue (and therefore cofactors interaction) did not play a crucial role for the activity of SRF fusions. This conclusion was also supported by the lack of effect of trametinib, a MEK1/2 inhibitor, on fusion activity (Fig. 2C-D), suggesting independence from MEK/ERK signaling, by contrast with the PHIP-BRAF fusion, used as a control in this experiment [22]. Expression of all constructs was verified by western blot (Supplementary Fig. S3A).

**Figure 2.**
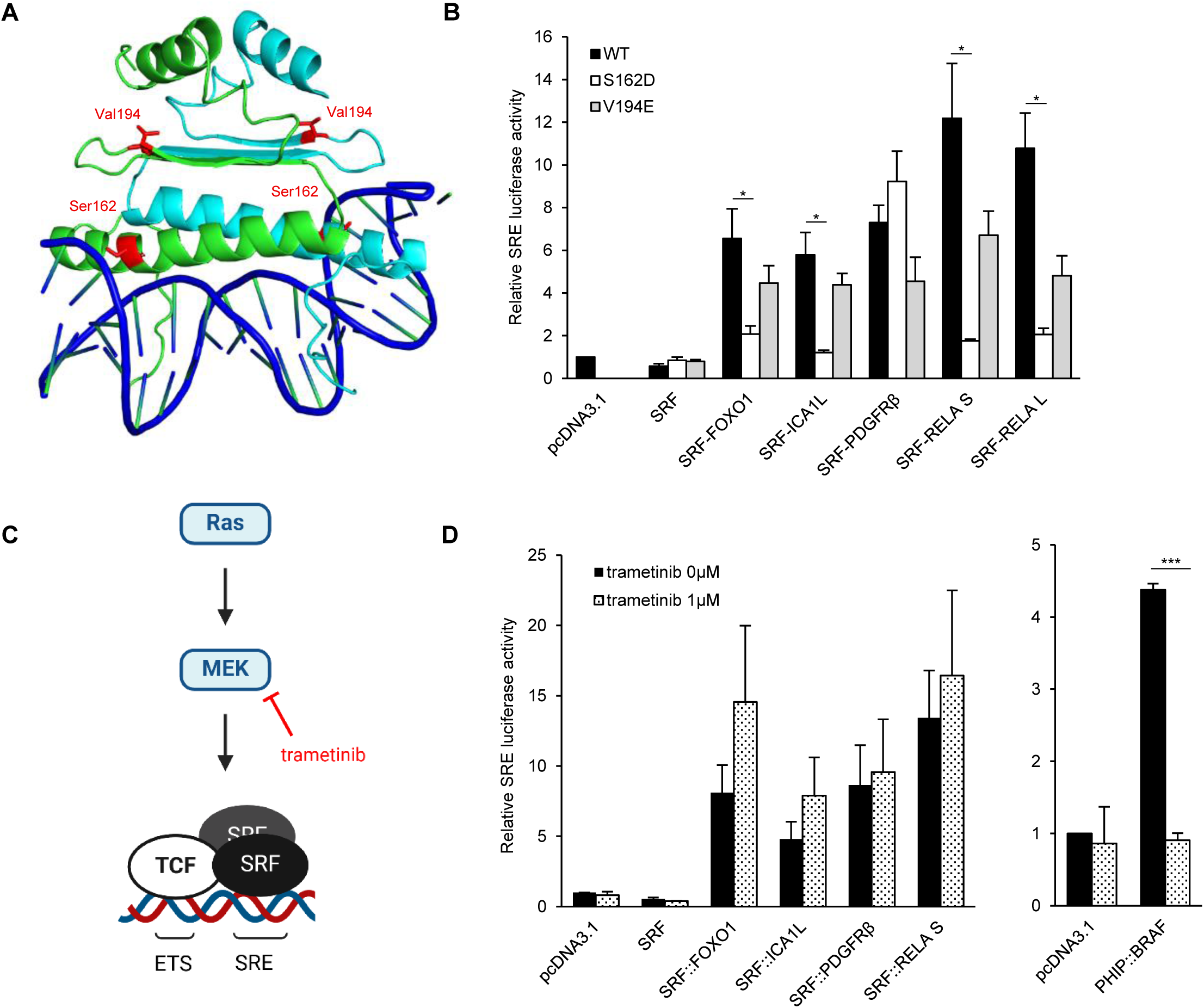
Role of SRF cofactors and MADS box in the transcriptional activity of fusions. (A) The 3D structure of the complex of the SRF MADS box with DNA was visualized using PyMOL (PDB ID: 1SRS). The DNA helix is represented in dark blue, and the two chains of the dimer of the MADS box in green and light blue. Amino acids of interest (Ser162 and Val194) are highlighted in red. (B) C2C12 were co-transfected with SRF fusions with or without mutation, SRE-luciferase reporter vector and an internal control. After 24h of transfection, luminescence was measured in cell lysates (N=4). (C) Schematic activation pathways of TCF signaling. (D) C2C12 were co-transfected with SRF fusions, SRE-luciferase reporter vector and an internal control. After 4h of transfection, cells were treated with 0 µM or 1 µM trametinib. After 24h of treatment, luminescence was measured in cell lysates (N=3). The mean of three independent experiments is represented with SEM (t-test with *p < 0.05; **p < 0.01; ***p < 0.001).

Together, these findings suggested that SRF fusions activity operated independently of TCF and MRTF cofactors but required DNA binding through the SRF MADS box, except for SRF::PDGFRβ.

### Most SRF fusions include a C-terminal transactivation domain

To further define the role of the 3’ partner for the fusion activity, we performed SRE luciferase assays after deletions or mutations of key domains and verified their expression by western blot (Supplementary Fig. S3B). Given that FOXO1 and RELA are established transcription factors, we first deleted their C-terminal TAD (the last 155 amino acids for FOXO1, and the last 101 amino acids for RELA) by introducing a stop codon in the sequence [23, 24]. As expected, TAD deletions suppressed the activity of SRF::FOXO1, SRF::RELA^L^ and SRF::RELA^S^ (Fig. 3A).

**Figure 3.**
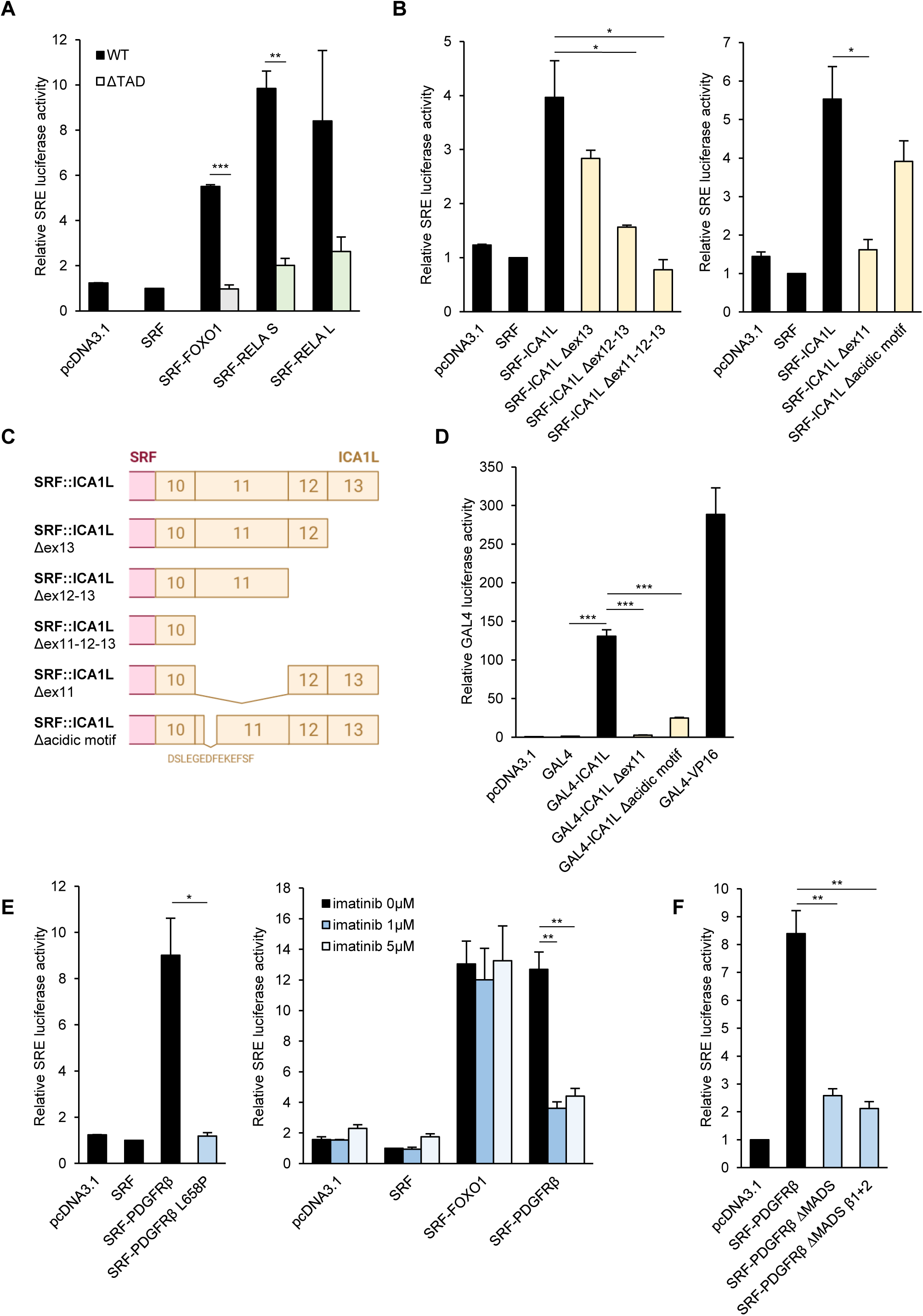
Role of C-terminal partners in the transcriptional activity of fusions. (A-B) C2C12 were co-transfected with SRF fusions with or without mutation, SRE-luciferase reporter vector and an internal control. After 24h of transfection, luminescence was measured in cell lysates (N=3). The TAD deletion corresponds to the TAD schematized in Figure 1A. (C) Schematic structures of SRF::ICA1L constructions. (D) HEK293T were co-transfected with GAL4 fused or not with the C-terminus of ICA1L (exons 10-13), pGL-TK-GAL4-luciferase reporter vector and an internal control. After 24h of transfection, the luminescence and the β-galactosidase activity were measured in cell lysates (N=3). (E-F) C2C12 were co-transfected with SRF fusions with or without mutation, SRE-luciferase reporter vector and an internal control. After 4h of transfection, cells were treated with 0 µM, 1 µM or 5 µM imatinib. After 24h of treatment, luminescence was measured in cell lysates (N=3). The mean of three independent experiments is represented with SEM (One-way ANOVA test with *p < 0.05; **p < 0.01; ***p < 0.001).

Next, we investigated whether the C-terminal region of ICA1L, a poorly characterized cytosolic protein, similarly influenced the transcriptional activity of the corresponding fusion. We sequentially deleted exons 13, 12 and 11 of *ICA1L* and observed a progressive decrease of the transcriptional activity, indicating that exons 11, 12 and 13 contributed to transcription (Fig. 3B-C). Most TADs are acidic-rich domains, such as the one found in RELA. Moreover, phenylalanine residues are crucial for the RELA TAD conformation, allowing interactions with the transcription preinitiation complex [24]. Exon 11 of *ICA1L* encodes a particularly high number of aspartates and glutamates, flanked by phenylalanine residues. Accordingly, exon 11 deletion led to a complete loss of luciferase signal, while removing a 14-amino-acid acidic motif (DSLEGEDFEKEFSF) at the beginning of exon 11 led to a partial loss of activity (Fig. 3B ; right panel). We confirmed these results using a GAL4 reporter assay in HEK293T cells, in which the C-terminal part of ICA1L (ex10-13) fused to the GAL4 DNA-binding domain strongly activated the luciferase reporter, supporting the unexpected transcriptional potential of ICA1L. Deletion of exon 11 or the 14-amino-acid acidic motif within exon 11 drastically suppressed the activity (Fig. 3D). The GAL4::VP16 fusion was used as a positive control. To explore whether the C-terminal tail of the SRF::PDGFRβ fusion, which is also rich in acidic residues, could harbor a cryptic TAD, we performed a similar luciferase assay (Supplementary Fig. S4A-B). However, no significant transcriptional activation was observed, indicating that this region does not act as a cryptic TAD.

Instead, the SRF::PDGFRβ fusion contains the intracellular part of the PDGFRβ receptor, including its tyrosine kinase domain. To determine the importance of this domain for the fusion activity, we introduced the PDGFRβ L658P mutation, previously described as a kinase dead variant, on the corresponding leucine in the fusion [25]. The SRE luciferase assay revealed a total loss of signal with L658P mutation (Fig. 3E ; left panel). Accordingly, imatinib (a tyrosine kinase inhibitor) significantly reduced SRF::PDGFRβ activity (Fig. 3E ; right panel).

Enforced dimerization plays a key role in the activation of PDGFRβ fusion proteins associated hematological malignancies [26, 27]. To determine if the MADS box was involved in dimerization and kinase transactivation, we deleted the entire MADS box (ΔMADS) or specifically its β1 and β2 strands (ΔMADS β1+2), key elements of the dimerization interface [28]. Both deletions significantly reduced transcriptional activity (Fig. 3F), indicating that the MADS box of SRF is critical for SRF::PDGFRβ function, alongside the tyrosine kinase domain.

In conclusion, these data highlighted the importance of C-terminal partners in the activity of SRF fusions: SRF::FOXO1, SRF::RELA and SRF::ICA1L acted as transcriptional activators via a canonical or cryptic TAD, while SRF::PDGFRβ relied on its kinase activity.

### SRF fusions undergo distinct dimerization patterns and compete with SRF WT on DNA

Since the MADS box is known to form dimers on DNA (Fig. 2A), we next tested the ability of SRF fusions to form homodimers or heterodimers with SRF WT, as tumor cells retained one intact SRF allele. Data from figures 2B and 3F already suggested that SRF::PDGFRβ activity may require dimerization without DNA binding. We performed co-immunoprecipitation (co-IP) experiments using Myc- and HA-tagged versions of SRF::RELA^S^ and SRF::PDGFRβ fusions. Western blot analysis revealed that Myc-SRF co-immunoprecipitated with HA-SRF, confirming a strong homodimerization of SRF WT. Both SRF::RELA^S^ and SRF::PDGFRβ were able to dimerize, forming homodimers as well as heterodimers with SRF WT. However, SRF::RELA^S^ preferentially formed heterodimers with SRF WT, whereas SRF::PDGFRβ showed a stronger tendency towards homodimerization (Fig. 4A).

**Figure 4.**
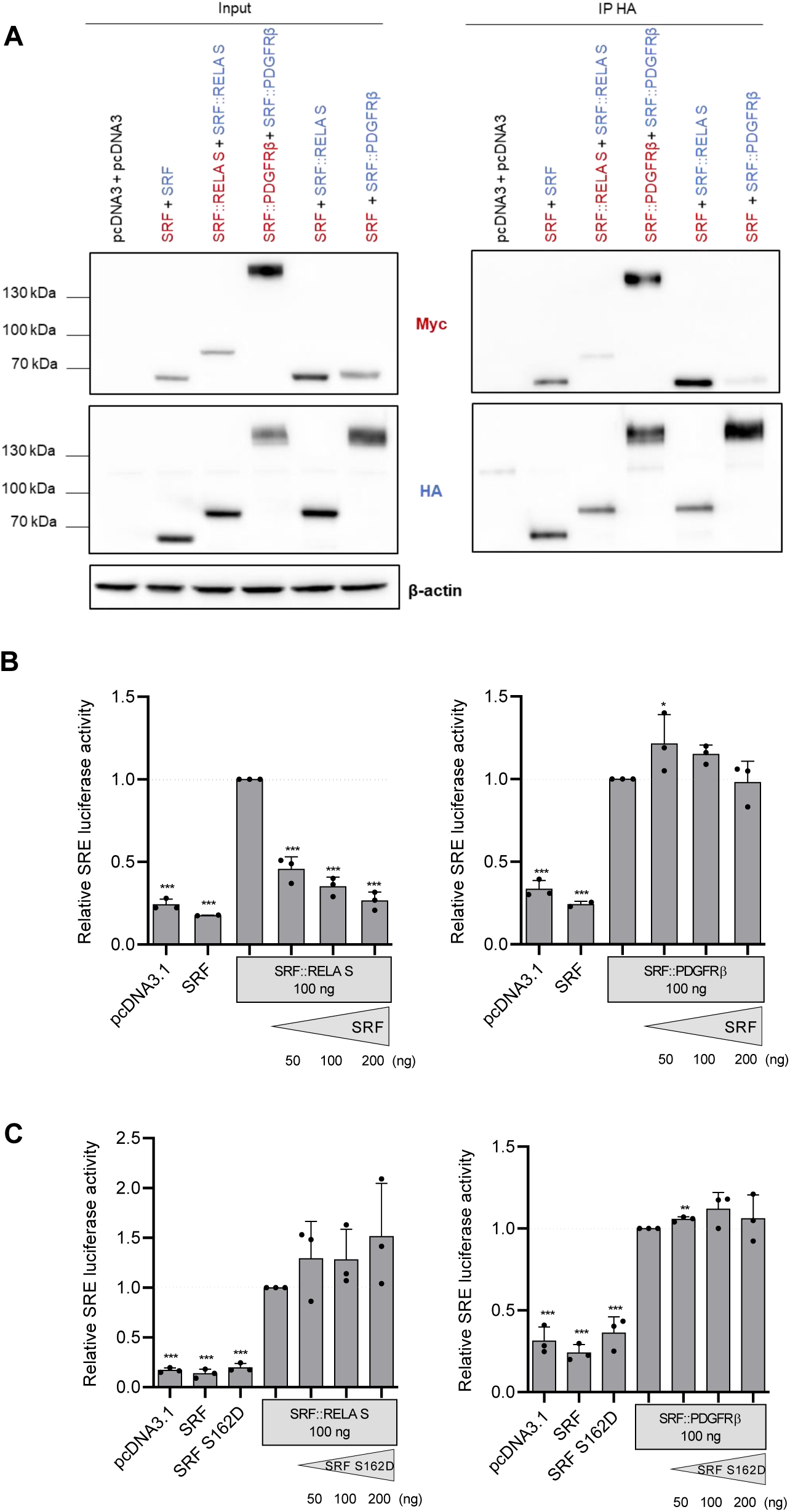
Dimerization mechanism of SRF fusions. (A) HEK293T cells were co-transfected with HA- and/or Myc-tagged SRF WT or fusions before protein extraction. SRF WT or fusions were immunoprecipitated with anti-HA antibody and analyzed by western blot with anti-HA and anti-Myc antibodies (N=3). (B,C) HEK293T were co-transfected with SRF-fusions and increasing amount of SRF WT or S162D (50, 100, 200 ng), pSRE-luciferase reporter vector and an internal control. After 24h of transfection, the luminescence and the β-galactosidase activity were measured in cell lysate (N=3) and normalized to SRF::RELA or SRF::PDGFRβ. The mean of three independent experiments is represented with SEM (One-way ANOVA test with *p < 0.05; **p < 0.01; ***p < 0.001).

We also tested the impact of SRF S162D mutation (to prevent DNA binding), which had no effect on protein-protein interaction, suggesting that MADS box dimerization occurs independently of DNA binding (Supplementary Fig. S5).

The influence of SRF WT on fusion activity was evaluated by SRE luciferase assays. Adding increasing amounts of SRF WT plasmid reduced transcriptional activity of SRF::RELA^S^ but not SRF::PDGFRβ (Fig. 4B). Adding increasing amounts of SRF S162D plasmid did not alter activity in either case, indicating that SRF WT competes with SRF::RELA^S^ for DNA binding (Fig. 4C).

In conclusion, SRF fusions exhibited distinct dimerization patterns: SRF::RELA^S^ interacted predominantly with SRF WT through heterodimerization, while SRF::PDGFRβ preferentially formed homodimers.

### SRF fusions promote cell growth

The ability of SRF fusions to enhance transcriptional activity suggested a potential role in oncogenic processes. We first examined their impact on the cell growth of mesenchymal stem cells (MSC), a relevant model given that SRF fusion-positive tumors exhibit mesenchymal and muscle-like features, although their precise cellular origin remains unclear. Since cells can quickly adapt to the expression of oncogenes, we used a doxycycline-inducible system to tightly control the expression of SRF fusions in MSC (Fig. 5A), enabling us to monitor cell growth after induction and reduce long-term adaptation effects. After three days of doxycycline treatment, cells expressing SRF fusions exhibited a modest yet significant increase in cell growth compared to non-induced cells and those expressing SRF WT (Fig. 5B). This effect was observed for all SRF fusions except SRF::PDGFRβ, which, in contrast, led to a pronounced loss of viability. This suggested that while most SRF fusions promote cell growth, SRF::PDGFRβ may engage distinct oncogenic mechanisms.

**Figure 5.**
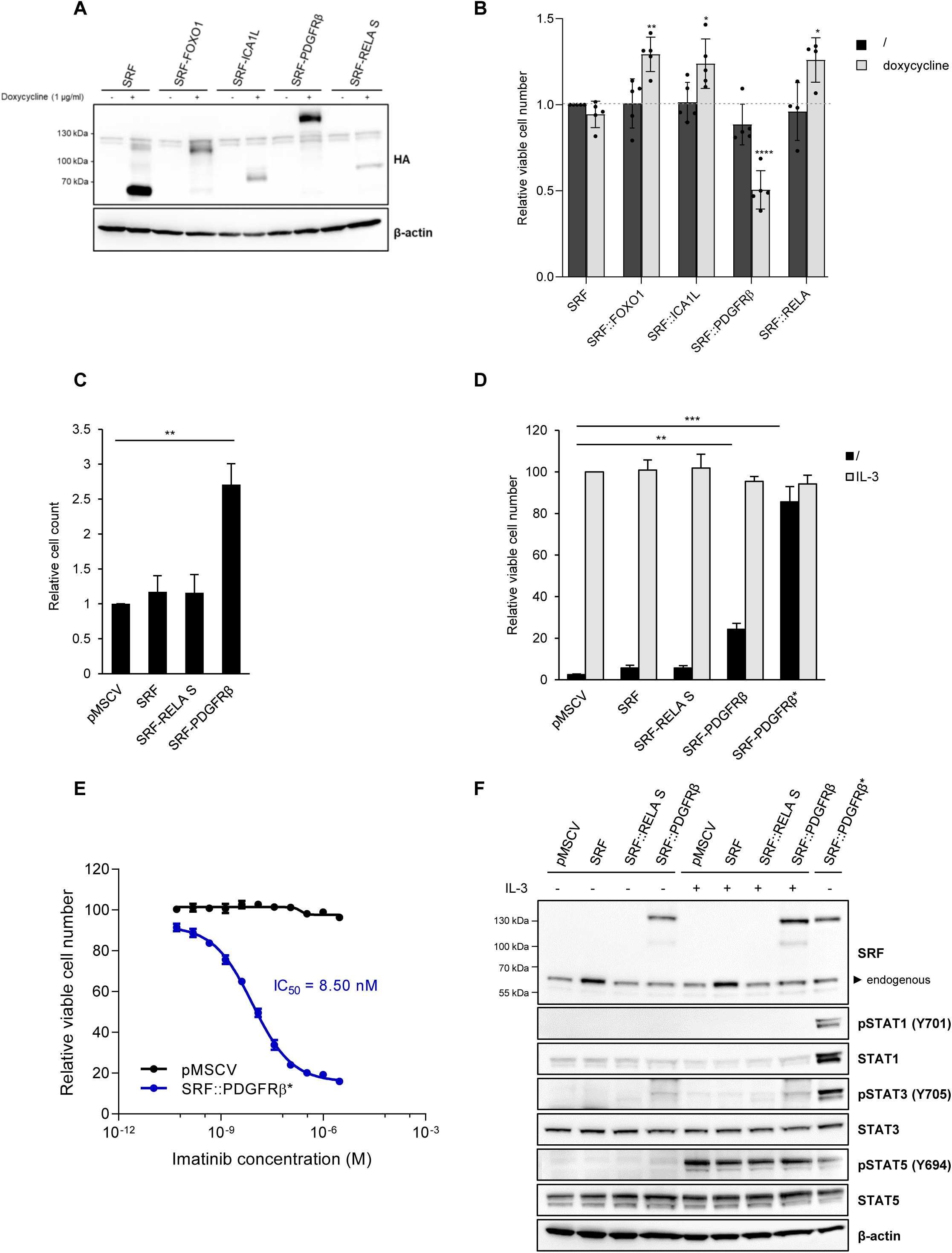
Most SRF fusions enhance cell growth in MSC, while SRF::PDGFRβ drives cytokine-independent survival in Ba/F3 cells. (A) MSC were transduced with pINDUCER-SRF fusions, and protein expression was analyzed by western blot after 24h of doxycycline treatment at 1 µg/ml (N=3). (B) CellTiter-Glo Luminescent assay was performed after 72h of doxycycline treatment at 1 µg/ml (N=5). (C) Ba/F3 cells were transduced with pMSCV-SRF fusions, and viable cells were counted 72h after IL-3 withdrawal (N=3). (D) CellTiter-Glo Luminescent assay was performed with or without IL-3 after 48h (N=3). SRF::PDGFRβ* grew independently of IL-3 for minimum 2 weeks. (E) SRF::PDGFRβ* and control cells were treated with imatinib, and IC50 was determined (N=3). (F) Protein expression and STAT phosphorylation were analyzed by western blot (N=3). The mean of three or more independent experiments is represented with SEM (Two-way ANOVA test or one-way ANOVA test with *p < 0.05; **p < 0.01; ***p < 0.001).

To further investigate whether SRF::PDGFRβ could favor proliferation in another cellular model, we stably expressed SRF::PDGFRβ in Ba/F3 cells, a widely used model for studying cellular transformation by oncogenic kinases [29]. In the presence of interleukin-3 (IL-3) in the culture media, a growth factor essential for Ba/F3 cell survival, no difference in proliferation was observed. In the absence of IL-3, cell viability was affected except for Ba/F3 cells expressing SRF::PDGFRβ (Fig. 5C-D). Cells expressing SRF::PDGFRβ were then adapted to culture in an IL-3-free medium, refer to as SRF::PDGFRβ*.

Subsequently, we assessed the sensitivity of these cells to imatinib (Fig. 5E). The half-maximal inhibitory concentration (IC50) was 8.50 nM, which is consistent with previously reported values for ETV6::PDGFRβ [27]. Since Ba/F3 cells survival relies on the activation of STAT transcription factors [30, 31], we explored the signaling mechanisms underlying the oncogenic potential of SRF::PDGFRβ. We observed constitutive phosphorylation of STAT1 and STAT3 in SRF::PDGFRβ* cells (Fig. 5F). In addition to Ba/F3 cells, STAT1 and STAT3 phosphorylation by SRF::PDGFRβ was observed in both MSC and C2C12 cells (Supplementary Fig. S6).

In conclusion, SRF::FOXO1, SRF::ICA1L and SRF::RELA^S^ fusions increased cell growth, while SRF::PDGFRβ enabled Ba/F3 cells to survive and proliferate in absence of IL-3. This ability, combined with the activation of STAT1 and STAT3 signaling pathways, suggests that SRF::PDGFRβ operates through distinct oncogenic pathways compared to other SRF fusions.

### Constitutive STAT activation requires SRF::PDGFRβ juxtamembrane domain truncation and dimerization

We next investigated in HEK293T the mechanism whereby SRF::PDGFRβ activates STAT1/3. Although enforced dimerization is a common mechanism of activation of *PDGFRB* fusions [26], we observed that the deletion of the MADS box or its β1 and β2-sheets had little impact on STAT1 and STAT3 phosphorylation (Fig. 6A-B). Therefore, we explored whether another mechanism could explain the constitutive activation of this pathway. Because of the fusion breakpoint, SRF::PDGFRβ lacks the 11 first amino acids of *PDGFRB* exon 12, truncating its auto-inhibitory juxtamembrane (JM) domain (Fig. 6A). Indeed, the rescue of a full JM domain decreased STAT1/3 phosphorylation. More striking decreases were observed when combined with deletions of the entire MADS box or its β-sheets (Fig. 6B ; Supplementary Fig. S7A). Interestingly, the phosphorylation of PDGFRβ itself was not impacted, at least on tyrosine 857.

**Figure 6.**
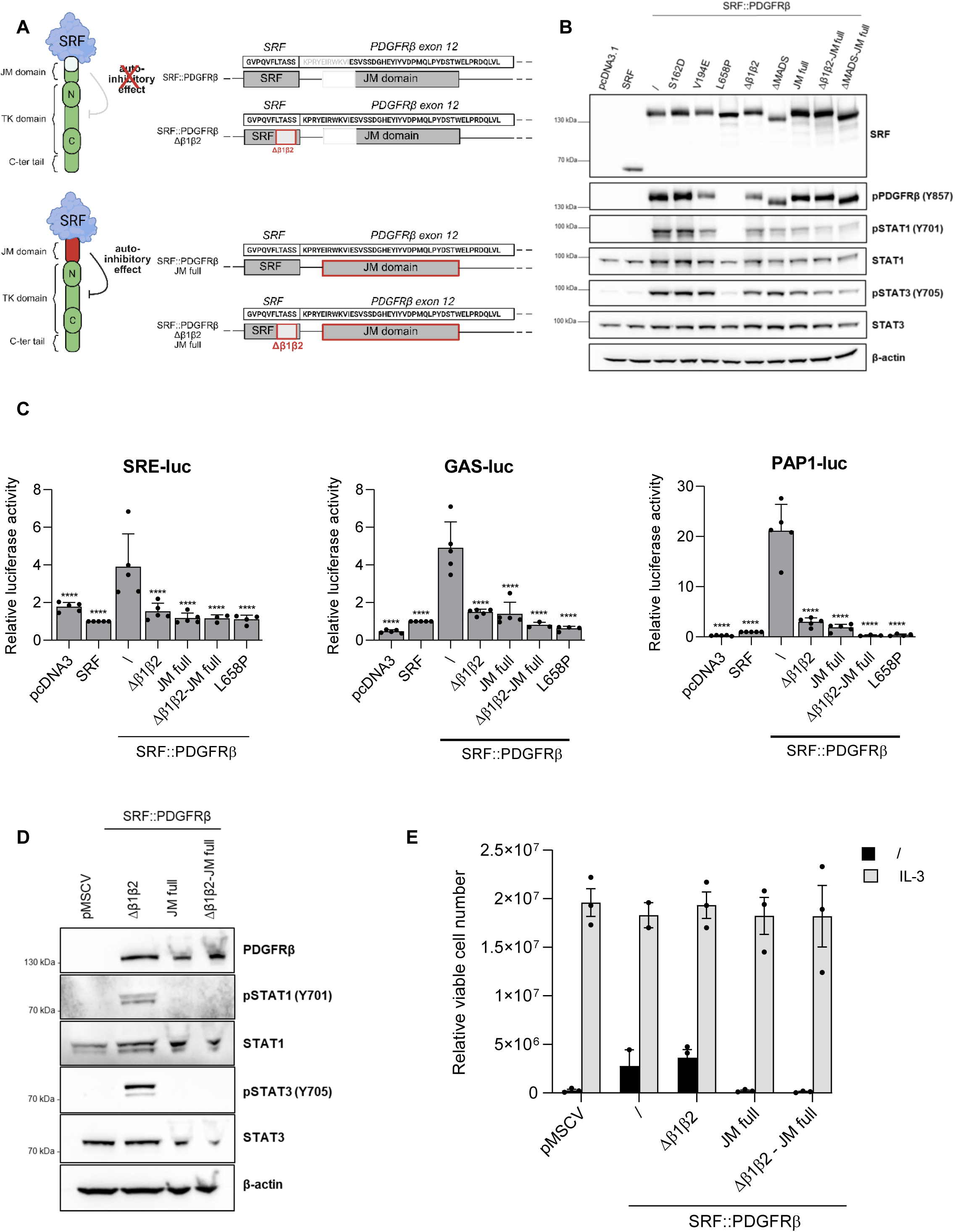
SRF::PDGFRβ activity relies on dimerization and JM domain truncation. (A) Schematic structures of SRF::PDGFRβ constructions. (B) Protein expression and STAT phosphorylation in HEK293T were analyzed by western blot (N=3). Quantification is available in Supplementary Figure 7A. (C) HEK293T were co-transfected with SRF fusions, luciferase reporter vectors (pSRE-luc for SRF activity, pGAS-luc for STAT1-mediated interferon response or pPAP1-luc for AP-1 activity downstream STAT3 pathway), and an internal control. After 24h of transfection, the luminescence and the β-galactosidase activity were measured in cell lysates (N=3). (D) Protein expression and STAT phosphorylation in Ba/F3 were analyzed by western blot (N=3). (E) CellTiter-Glo Luminescent assay was performed in the presence or absence of IL-3 after 48h of culture (N=3). The mean of three independent experiments is represented with SEM (One-way ANOVA test with *p < 0.05; **p < 0.01; ***p < 0.001; ****p < 0.0001).

In line with these results, luciferase reporter assays associated with various SRF- and STAT-driven promoters showed that deleting the β-sheets or restoring the full JM domain significantly reduced transcriptional activity, and combining both deletions abolished activity entirely (Fig. 6C ; Supplementary Fig. S7B). The kinase dead mutant (L658P) was used as a control. Finally, in Ba/F3 cells, partial truncation of the JM domain is sufficient to autoactivate the kinase, as shown by the loss of STAT1 and STAT3 phosphorylation when the domain is intact (Fig. 6D). Moreover, the oncogenic effect of SRF::PDGFRβ in Ba/F3 cells was lost with a full JM domain, leading to cell death in absence of IL-3 (Fig. 6E).

These findings demonstrated that the oncogenic activity of SRF::PDGFRβ depends on the partial deletion of the auto-inhibitory juxtamembrane domain of PDGFRβ, as well as MADS-driven dimerization.

### SRF fusions induce specific gene expression profiles related to muscle development and inflammation

The aberrant transcriptional activity and proliferative advantage conferred by SRF fusions suggested that they reshaped transcriptional programs to sustain oncogenic properties. Therefore, we analyzed their impact on global gene expression by performing RNA-sequencing on both MSC and C2C12 lines stably expressing SRF fusions (Fig. 7 ; Supplementary Fig. S8-9). Differential gene expression analysis (normalized on empty vector conditions) revealed that all four fusions induced significant transcriptional changes compared to SRF WT, with large numbers of down- and upregulated genes across both models (Fig. 7A). SRF::FOXO1, SRF::ICA1L and SRF::RELA share a substantial number of regulated transcripts in MSC (n=124) and in C2C12 (n=175), suggesting that they could operate through overlapping mechanism [13, 16]. By contrast, SRF::PDGFRβ shared fewer regulated genes (Fig. 7A). As expected by their increased SRE luciferase activities, all fusions (except SRF::PDGFRβ in C2C12) maintained or even enhanced the expression of canonical SRF target genes (Supplementary Fig. S8; SRF/MRTF gene set from [32]). Notably, SRF fusions significantly modulated the expression of genes associated with the myogenesis hallmark in both models, but in opposite directions (Fig. 7B). In MSC, genes such *ACTA1, ACTC1* and *IGFBP3* were strongly upregulated, suggesting forced or accelerated differentiation of MSC into muscle-like cells. This observation was supported by the significant enrichment of the biological process “muscle structure development” in all four conditions (Supplementary Fig. S9A). In contrast, in committed myogenic C2C12 myoblasts, key differentiation markers such as *Myog*, *Myl1* and *Des* were downregulated, consistent with an impairment of terminal differentiation (Fig. 7B). Some of the differentially expressed genes were validated by RT-qPCR in MSC (Supplementary Fig. S10).

**Figure 7.**
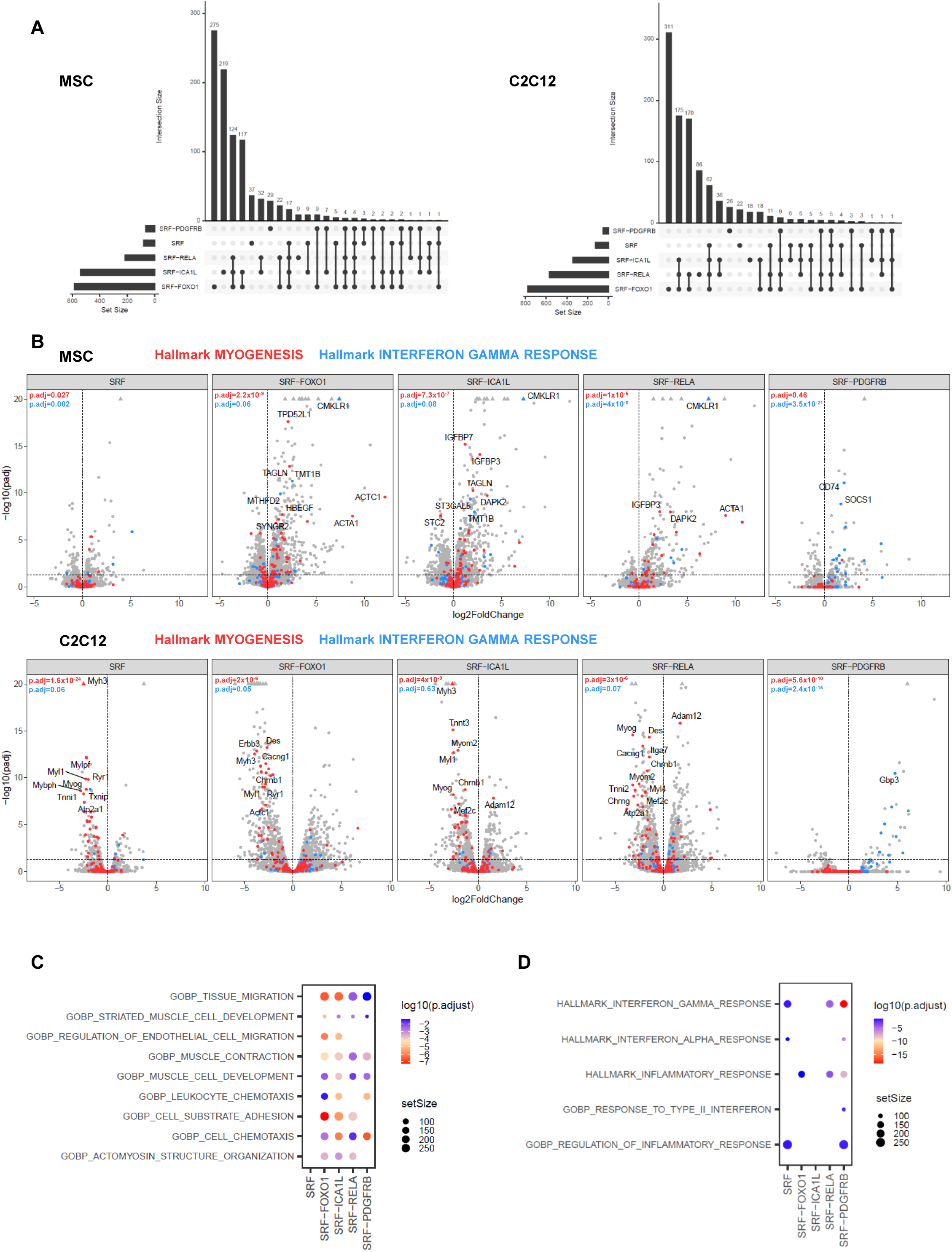
SRF fusions and gene expression profiles in C2C12 and MSC. (A) Visualization of “hallmark myogenesis” (labelled in red) and “hallmark interferon gamma response” (labelled in blue) gene sets for SRF, SRF::FOXO1, SRF::ICA1L, SRF::RELA, and SRF::PDGFRβ conditions. (B) Comparison of differentially regulated genes between SRF, SRF::FOXO1, SRF::ICA1L, SRF::PDGFRβ, SRF::RELA S. Gene sets comprise significantly up- and downregulated genes compared to empty control vector pMSCV (log2 fold change > 1; adjusted p-value > 0.5). (C) GO Biological Process (BP) enrichment analysis of genes upregulated in MSC expressing SRF fusions. For each fusion, we selected the top 10 most significantly enriched GO Biological Process terms (based on adjusted p-value) and removed redundant or overlapping terms. (D) Enrichment of inflammation-related gene sets in MSC expressing SRF fusions.

Despite overall similarities, the fusions displayed distinct transcriptional outputs, especially SRF::PDGFRβ, which showed robust and specific upregulation of interferon response genes, particularly within the IFN-γ pathway (Fig. 7B-D ; Supplementary Fig. S9B). This included upregulation of chemokines, such as *CXCL9*, *CXCL10* and *CXCL11*, consistent with the activation of the STAT1 pathway by SRF::PDGFRβ, and its association with inflammatory myofibroblastic tumors.

To further explore the biological processes activated by these fusions, we performed a gene set enrichment analysis (GSEA) and visualized the top 10 significantly upregulated biological processes in MSC (Fig. 7C). All four fusions strongly induced muscle-related processes. Importantly, SRF::PDGFRβ again stood out by selectively enriching for immune and interferon-related pathways (Fig. 7D), supporting its distinct transcriptional output.

In conclusion, while all SRF fusions drive expression of genes associated with muscle development, SRF::PDGFRβ uniquely activates inflammatory pathways.

### Strong enrichment of patients SRF-rearranged perivascular myoid tumors gene signature in the transcriptome of MSC expressing SRF fusions

To assess whether the transcriptional program driven by SRF fusions in MSC mirrors the gene expression landscape of human tumors, we evaluated the enrichment of genes upregulated in SRF fusion-positive tumors within our MSC models. We first performed a GSEA using a published dataset of 13 soft tissue tumors harboring SRF fusions [16]. This analysis revealed a significant enrichment of the upregulated tumor gene signature across MSC lines expressing SRF::FOXO1, SRF::ICA1L and SRF::RELA S (data not shown). To strengthen this observation, we extended the analysis to a larger cohort of 35 SRF-rearranged perivascular myoid tumors [10, 12, 13, 16]. Consistent with our initial results, the transcriptome of MSC expressing SRF fusions showed a highly significant enrichment for genes upregulated in SRF fusion-positive tumors (Fig. 8 ; Supplementary Fig. S11). This strong concordance confirmed that MSC models reproduced the transcriptional effects of SRF fusions observed in patient tumor samples. Many of shared up-regulated genes between MSC and human tumors were involved in muscle differentiation, including *ACTG2, MYH11, CNN1, LMOD1* and *CDH3*. Several of these genes were independently validated by RT-qPCR in MSC lines, with log₂ fold changes up to 8 (Supplementary Fig. S10).

**Figure 8.**
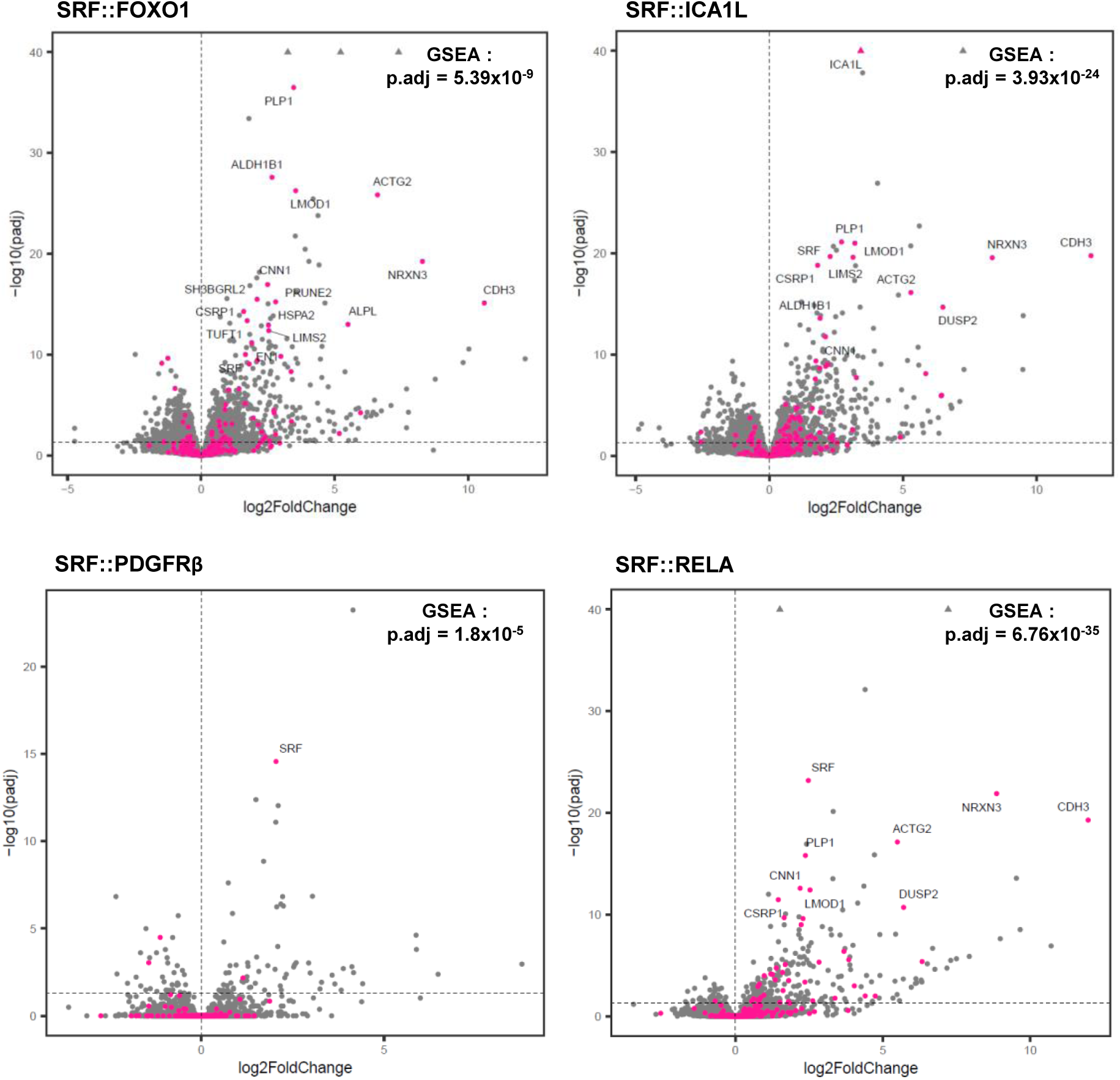
Genes upregulated in SRF fusion–positive tumors are enriched across MSC models expressing distinct SRF fusions. Volcano plots showing differential gene expression in MSC expressing SRF fusions compared to empty control vector pMSCV. Genes identified as significantly upregulated in SRF-rearranged perivascular myoid tumors (n=35) relative to fusion-negative myofibromas (n=38), based on a Bonferroni-adjusted p-value < 1×10⁻⁵ and a fold change ≥ 3, are highlighted in pink. Adjusted p-values from GSEA confirm their significant enrichment in the MSC models.

Together, these results demonstrate that SRF fusions impose a muscle-like transcriptional identity on MSC, mimicking the molecular signature observed in human tumors.

## DISCUSSION

Although the SRF transcription factor has been thoroughly studied for decades, SRF gene alterations in cancer were discovered only recently. In the present study, we demonstrate that *SRF* fusion genes encode nuclear oncoproteins associated with a constitutive transcriptional activity, unlike wild-type SRF, which regulates gene expression in response to extracellular signals with the tightly regulation of specific cofactors. Our results suggest that SRF fusions bind directly to the promoter of SRF target genes, and activate transcription through the acquired partner TAD, which alleviates the need for SRF cofactors recruitment and may even facilitate interactions with alternative factors, thus broadening their transcriptional potential.

SRF fusions were exclusively found in soft tissue tumors expressing muscle differentiation markers. In line with this observation, RNA-sequencing performed on mesenchymal stem cells expressing the different SRF fusions revealed a striking upregulation of smooth and skeletal muscle markers, including *ACTA1, ACTA2*, *ACTC1, MYH11, TAGLN* or *CNN1*. Gene ontology analyses consistently highlighted biological processes related to muscle development and differentiation. This simultaneous activation of distinct myogenic programs occurred without external stimulation, indicating an intrinsic deregulated transcriptional state rather than a physiologically-induced differentiation process. The resulting myoid transcriptional signature reflects the phenotype observed in human tumors and supports a direct role for SRF fusions in promoting myogenic gene expression in mesenchymal cells. Supporting this hypothesis, the transcriptional profiles of MSC expressing SRF fusions show a strong enrichment for genes upregulated in SRF fusions-positive tumors, highlighting the relevance of these models for investigating SRF-driven tumorigenesis.

Beyond muscle-related genes, SRF fusions also aberrantly activate unrelated genes such as *Plp1* and *Nrxn3*, which are involved in the nervous system, suggesting broader transcriptional dysregulation. The presence of non-native transactivation domains in the fusions may alter chromatin accessibility or redirect cofactor recruitment, leading to inappropriate gene expression and potentially contributing to oncogenic transformation. Similar mechanisms have been described for other fusion-driven cancers. For example, in alveolar rhabdomyosarcoma, PAX3::FOXO1 establishes myogenic super-enhancers that drive aberrant gene activation and create a dependency on BET chromatin regulators [33]. Similarly, the EWS::FLI1 fusion in Ewing sarcoma reprograms gene regulatory circuits by directly modulating enhancer activity [34].

Our study allows the classification of C-terminal fusion partners into three categories. In the first one, SRF is fused to a TAD of another known transcriptional regulator, such as RELA and FOXO1, creating new hybrid transcription factors with the MADS box of SRF and the TAD of a partner. Both tested fusions stimulated MSC growth to some extent. The mechanism is reminiscent of PAX3::FOXO1 in rhabdomyosarcoma, where the fusion of a DBD to a strong TAD hijacks developmental transcriptional programs. This mechanism of activation may also be valid for SRF::E2F1, SRF::STAT6, SRF::MKL2, SRF::NCOA1/2/3, and SRF::CITED1. As expected, the DNA-binding domain (RHD) of RELA, retained in SRF::RELA^L^ but not in SRF::RELA^S^, does not seem to impact SRE activity. However only SRF::RELA^L^ activated a NF-κB luciferase reporter. While the RELA RHD is essential for oncogenicity in supratentorial ependymomas [35], SRF::RELA^S^ still promoted cell growth.

A second category includes SRF fusions with a non-physiologic cryptic TAD, as seen with SRF::ICA1L. Although ICA1L is not a known transcriptional regulator, our data suggest that its C-terminal region harbors activation potential once forced into the nucleus by SRF fusion. Initially involved in vesicle trafficking, ICA1L becomes a potent transcriptional activator in this context, likely through acidic residues in exon 11 that mimic classical TAD. In the end, the fusion results in a functional unit combining SRF DBD with an unexpected TAD. Similarly, the C3orf62 protein was recently described to hold a TAD in its C-terminal region, to interact with CBP/p300 in the nucleus [36], and to activate SRE targets. These cases underscore how forced nuclear relocalization and association with SRF MADS box can reveal latent transactivation capacity in non-nuclear proteins.

The third group comprises SRF fusions devoid of any TAD, represented by the SRF::PDGFRβ fusion, for which we confirmed the absence of TAD activity. In this case, transcriptional activation is fully dependent on tyrosine kinase activity, as evidenced by the kinase dead mutant and the sensitivity to imatinib. The fusion retains a constitutively active kinase, due to a partial truncation of the auto-inhibitory juxtamembrane domain in exon 12 of *PDGFRB*, a mechanism also observed in FIP1L1::PDGFRα and engineered mutants of ETV6::PDGFRβ [37, 38]. This truncation likely relieves the auto-inhibition and enables kinase activation even in the absence of ligand or oligomerization. Our data suggest that SRF::PDGFRβ activation also depends on dimerization via the MADS box of SRF, particularly through its β-sheets, which are known to mediate SRF homodimerization. This mechanism mirrors the role of the helix-loop-helix (HLH) domain in ETV6::PDGFRβ, which is responsible for the fusion oligomerization and the subsequent activation of the kinase [38]. Altogether, SRF::PDGFRβ appears to use a dual mechanism combining constitutive kinase activity and dimerization to drive transcriptional activation in the absence of a TAD. In addition to direct or indirect STAT phosphorylation, we hypothesize that SRF::PDGFRβ activates the remaining endogenous SRF WT indirectly, which would explain the specific SRE activity and gene expression profile. Functionally, SRF::PDGFRβ shows a distinct oncogenic profile compared to other SRF fusions by stimulating the growth of Ba/F3 cells, but not MSC. We speculate that the overactivated PDGFR signaling may lead to strong STAT1 activation, known to mediate anti-proliferative responses and interferon-induced cell death [39]. Accordingly, SRF::PDGFRβ induces an interferon-like transcriptional program in both C2C12 and MSC. SRF::PDGFRβ was identified in a case of inflammatory myofibroblastic tumor, raising the possibility that this IFN-like response may contribute to the associated immune infiltrate. However, activating point mutations in PDGFRβ are more frequent in ‘non-inflammatory’ myofibromas, which belong to the perivascular tumor family, like SRF-fusion positive tumors. The particular nuclear localization of SRF::PDGFRβ compared to activating mutations affecting the transmembrane PDGFRβ could explain the different oncogenic mechanism and tumor phenotypes. Indeed, all SRF fusions were detected in the nucleus, regardless of the nature of the C-terminal partner. They all conserved the nuclear localization signal (NLS) of SRF, located in its first exon [40]. But the functional impact of the nuclear localization of SRF::PDGFRβ remains unclear.

Taken together, our study demonstrates that SRF fusion genes are oncogenes that drive constitutive transcriptional activity and promote cell growth. Despite various fusion partners, they converge to the aberrant activation of SRF target genes and induce a myoid transcriptional program, consistent with the phenotype of SRF fusion-positive human tumors. While SRF::PDGFRβ could be blocked by tyrosine kinase inhibitors such as imatinib, further studies are needed to explore therapeutic strategies to target SRF fusions that encode chimeric transcription factors.

## MATERIALS AND METHODS

### Cell culture, reagents and vectors

Immortalized human mesenchymal stem cells (MSC-hTERT were cultured in Minimum Essential medium α (MEM α) (Gibco, Watham, MA, USA) supplemented with 10% HyClone serum, 1% L-glutamine, 1 ng/ml FGF-2, 0.2 mM ascorbic acid, 2 U/ml heparin and antibiotics (50 U/ml penicillin and 50 μg/ml streptomycin). Mouse C2C12 myoblasts, a kind gift from Pr. D. Tyteca (de Duve Institute, Brussels, Belgium), and human embryonic kidney HEK293T cells (ATCC, Manassas, VA, USA) were cultured in Dulbecco’s modified Eagle’s medium (DMEM) (Lonza, Basel, Switzerland) supplemented with 10% fetal bovine serum. Ba/F3 cells were grown in the same medium supplemented with IL-3 (500 U/ml) [41]. All cells were maintained at 37 °C in 5% CO₂. Imatinib and trametinib were purchased from LC Laboratories (Woburn, MA, USA) and Selleckchem (Houston, TX, USA), respectively. Human coding sequences of SRF, SRF::FOXO1, SRF::ICA1L, SRF::PDGFRB, SRF::RELA^S^, and SRF::RELA^L^ were obtained from Genscript (Piscataway, NJ, USA). Exons 10-13 of ICA1L (C-terminus) were amplified by PCR and cloned into the pcDNA3.1-GAL4 vector using XbaI restriction sites. Mutations and deletions were introduced using the QuickChange XL-II kit (Agilent, Santa Clara, CA, USA) following the manufacturer’s protocol (see Supplementary Table 1). SRF fusions were subcloned into the retroviral pMSCV vector via NotI sites [42], and in into the lentiviral pINDUCER vector using the In-Fusion Cloning system (Takara, France). Expression from pINDUCER constructs was induced with 1 µg/ml doxycycline (#D9891, Sigma-Aldrich, Saint-Louis, MO, USA). All constructs were verified by Sanger sequencing (Eurofins, Luxembourg).

### Cell transfection

C2C12 and MSC cells were transiently transfected with Lipofectamine 2000 reagent (Invitrogen, Waltham, MA, USA) as described [43]. HEK293T cells were transfected with the calcium phosphate precipitate method as described [44].

### Retroviral and lentiviral infections

Stable C2C12, MSC and BaF/3 cell lines expressing SRF fusions were generated using the retroviral pMSCV vector. Viral particles were produced by HEK293T cells seeded in T75 flasks. pMSCV-SRF WT or fusion vector (18 µg) was co-transfected with the packaging plasmid pCMV-Gag-Pol (6 µg) and the envelope plasmid pCMV-VSVg (3 µg). Supernatants were collected 48 h after transfection and cells were infected in presence of polybrene (8 µg/ml). After 24 h, cells were washed and selected with puromycin (1 µg/ml) for 5 days. For MSC infection with pINDUCER, lentiviruses were generated with the same amount of transfected plasmids, but using pCMV-dR8.2 as the packaging plasmid. After infection, cells were sorted for GFP+ populations using the MA900 cell sorter (Sony Technology, USA).

### Protein extraction, western blot and co-immunoprecipitation

Transiently transfected or transduced cells were washed with PBS and lysed for 20 min on ice in lysis buffer (25 mM Tris-HCl pH 7.4, 150 mM NaCl, 6 mM EDTA, 10% glycerol, 1% Triton X-100) supplemented with protease inhibitors (1 mM sodium orthovanadate, 1 mM Pefabloc, 1 µg/ml aprotinin). Lysates were cleared by centrifugation (10 min, 4°C), and protein concentration was determined using the BCA Protein Assay Kit (Thermo Fisher Scientific, USA). Samples (30 µg) were mixed with 4x Laemmli buffer (0.2 M Tris-HCl pH 6.8, 8% SDS, 0.4% bromophenol blue, 40% glycerol, 2.8% β-mercaptoethanol), boiled for 5 min, and analyzed by western blot using Novex Tris-Glycine 4–12% gels (Thermo Fisher Scientific) and PVDF membranes (Amersham Hybond P, GE Healthcare). Membranes were blocked in 5% milk for 1 h at room temperature, then incubated overnight at 4°C with the indicated primary antibodies : anti-HA-tag (#3724, Cell Signaling Technology, Danvers, MA, USA), anti-Myc-Tag (#2272, Cell Signaling Technology), anti-phospho-STAT1 (#9167, Cell Signaling Technology), anti-STAT1 (#610115, BD BioSciences), anti-phospho-STAT3 (#9145, Cell Signaling Technology), anti-STAT3 (#12640, Cell Signaling Technology), anti-phospho-STAT5 (#9351, Cell Signaling Technology), anti-STAT5 (#94205, Cell Signaling Technology), anti-SRF (#5147, Cell Signaling Technology), anti-phospho-PDGFRβ (#3170, Cell Signaling Technology), anti-PDGFRB (#3169, Cell Signaling Technology), anti-β-actin (#A5441, Sigma-Aldrich). Detection was performed using chemiluminescence (Fusion Solo S, Vilber).

For co-immunoprecipitation, cells were lysed in RIPA buffer (50 mM Tris-HCl pH 7.4, 150 mM NaCl, 1 mM EDTA, 0.25% sodium deoxycholate, 1% NP-40) with protease inhibitors as above. Lysates (800 µg) were incubated overnight at 4°C with anti-HA or anti-Myc antibodies, followed by incubation with Protein A/G UltraLink Resin (#53133, Thermo Fisher) for 1.5 h at 4°C. After three washes, samples were resuspended in 2x Laemmli buffer for western blot analysis.

### Immunofluorescence

C2C12 cells were transfected on coverslips in 6-well plates (200 000 cells/well). After 2 successive washes with PBS, cells were incubated in fixation buffer (3% formaldehyde, 2% sucrose, diluted in PBS) for 15 min at room temperature. Cells were washed 3 times for 3 min with PBS and incubated in permeabilization buffer (20 mM Tris-HCl pH 8, 50 mM NaCl, 3 mM MgCl_2_, 10% sucrose, 0.5% Triton X-100) for 10 min at room temperature. Cells were washed 3 times for 3 min with PBS and blocked in 1% bovine albumin serum solution for 1 h at room temperature and incubated overnight at 4°C with the anti-HA-tag antibody (#3724, Cell Signaling Technology). The next day, cells were washed 3 times for 3 min with PBS-Tween20 0.1% and incubated with Hoechst 33258 (dilution 1/5000, Invitrogen), Texas Red X-phalloidin (dilution 1/100, Molecular Probes), and secondary antibody coupled with Alexa Fluor 488 fluorochrome (dilution 1/400) for 45 min at room temperature. Cells were finally washed 3 times for 3 min with PBS-Tween20 0.1% and 3 times for 3 min with PBS before mounting with Dako Fluorescence Mounting Medium (Agilent). Images were captured with the Cell Observer Spinning Disc Confocal microscope (Zeiss, Iena, Germany) and analyzed with the ImageJ Fiji software.

### Luciferase reporter assay

C2C12 cells were co-transfected with pcDNA3.1-SRF WT or fusion (150 ng), SRE-firefly luciferase reporter vector (150 ng) and pRL-TK renilla luciferase vector (150 ng) as internal control. In some cases, 4 h after transfection, cells were treated with imatinib or trametinib at the indicated concentrations for 24 h at 37°C. Firefly and renilla luciferase luminescences were measured in cell lysates using the Dual-Luciferase Reporter Assay System (Promega, Madison, WI, USA) as described [45]. For GAL4, GAS, PAP1 and SPI reporter assays, HEK293T cells were co-transfected with the expression vector of interest (pcDNA3.1, 150 ng), the luciferase reporter vector (pGL-TK-GAL4-luc, pGAS-luc, pPAP1-luc or pSPI-luc, 150 ng) and pEF-β-galactosidase (150 ng) as internal control, and firefly luciferase activity was monitored [44].

### CellTiter-Glo luminescent viability assay

Transduced Ba/F3 cells were washed three times with PBS to eliminate IL-3 from the medium. Cells were then seeded in 96-well plates (30 000 cells/well) in 200 µl of complete medium, with or without IL-3 in four replicates for each condition, and incubated for 48 h at 37°C. Inducible-MSC were seeded in 96-well plates (100 000 cells/well) in presence of doxycycline (1 µg/ml), and incubated for 72 h at 37°C. CellTiter-Glo reagent (100 µl, #G7572, Promega) was added to each well, followed by a 15 min incubation at room temperature. The luminescence was then measured using a GLOMAX instrument (Promega) as described [46].

### RNA extraction and RT-qPCR

Total RNA was extracted with TriPure Isolation Reagent (Sigma-Aldrich) and subjected to reverse transcription using M-MLV reverse transcriptase (Thermo Fisher Scientific). Quantitative PCR was performed using the KAPA SYBR® FAST qPCR Kit (Kapa Biosystems, Roche) according to the manufacturer’s instructions and fold-changes were calculated with the ΔΔCt method.

### RNA-sequencing and analysis

Total RNA of transduced MSC and C2C12 cell lines was extracted as mentioned above and cleaned from residual genomic DNA with DNase on column (NucleoSpin RNA, Macherey-Nagel, Düren, Germany). Libraries were prepared using the TruSeq Stranded mRNA kit and sequenced on a NovaSeq 6000 Sequencing System (Illumina, San Diego, CA, USA). The experiment was carried out in triplicates (duplicates for SRF::PDGFRβ in C2C12) on independent cell transductions. This generated 30 million paired-end reads of 150 nucleotides per specimen (Macrogen, Seoul, South Korea). Read quality control was performed using FastQC software v0.11.8 (Andrews S, Babraham Bioinformatics). Low quality reads were trimmed and adapters were removed using Trimmomatic software v0.39 [47]. Reads were aligned using the HISAT2 software v2.12.1 [48] on grch38 (MSC) or GRCm38 (C2C12) genome. Gene counts were generated using featureCounts software from subread-2.0.3 (MSC) or subread-2.0.0 (C2C12) [49] and Ensembl Homo_sapiens.GRCh38.105.gtf (MSC) or Mus_musculus.GRCm38.94.gtf (C2C12) annotation file. Differential expression analyses were done using DESeq2 v1.42 [50], on R v4.3.0. Gene set enrichment analyses were performed with clusterProfiler v4.10.0 bioconductor package.

For comparative transcriptomic analysis with MSC cell lines, we used RNA sequencing data from perivascular tumors. This study was approved by our local internal review board and patients gave their consent. Twenty SRF-rearranged perivascular myoid tumor and 38 myofibroma samples were identified and RNA-sequenced during molecular diagnostic routine in the Leon Bérard Cancer Center (Lyon, France). Fiftheen additional SRF-rearranged perivascular myoid tumor was provided MSKCC (New York, USA). Tumor samples RNA extraction, whole Exome RNA sequencing and expression profiles were performed as described [51]. Differentially expressed genes between SRF-rearranged perivascular myoid tumors (n=35) and myofibromas (n=38) were assessed by a Welsh t-test comparison (in the R environment) and considered significant if their absolute fold change were above 2 with a Bonferroni adjusted p-value below 0.01.

The significantly upregulated genes were selected based on Bonferroni-adjusted p-values < 1×10⁻⁵ and a fold change ≥ 3, yielding 340 upregulated genes. These genes were highlighted in the volcano plots generated from MSC cell lines expressing SRF fusions, and the resulting signature was subsequently tested for enrichment by Gene Set Enrichment Analysis (GSEA).

### Statistical analysis

All experiments were independently repeated at least three times. Data were normalized to SRF WT or empty vector unless otherwise specified, and are presented as mean ± standard error of the mean (SEM). Statistical analysis was performed using bilateral Student’s t-test for comparisons between two groups. For multiple group comparisons, one-way ANOVA or two-way ANOVA were used, with Brown-Forsythe and Welch corrections when appropriated. Statistical significance was denoted as follows: *, p<0.05; **, p<0.01; ***, p<0.001; ***, p<0.0001. Non-significant differences are not indicated.

## ACKNOWLEDGEMENTS

Constance Pirson and Ariane Sablon are the recipients of Aspirant fellowships from the Fonds de la Recherche Scientifique - FNRS, (grants #40017117 and 40000851, respectively). This work was also supported by FNRS grant #J.0062.22 (to J.B.D.). We are grateful to Stefan Constantinescu, Donatienne Tyteca and Paul Coffer for generous donation of reagents, and to Sandrine Lenglez for technical support.

## AUTHOR CONTRIBUTIONS

J-B.D., A.S. and C.P. conceptualized the experiments. C.P. and A.S. performed the experiments and analyzed the data. P.A. contributed to some experiments. A.L. carried out the bioinformatic analyses. A.K.B., M.K., C.R.A. and F.T. provided key samples and data. J-B.D. was responsible for funding acquisition and supervision. The original draft was written by J-B.D., C.P. and A.S., and all authors contributed to the review and editing of the manuscript.

## COMPETING INTERESTS

The authors declare no competing interests.

## DATA AVAILABILITY

RNA sequencing data generated in this study have been deposited in the Gene Expression Omnibus (GEO) under accession codes GSE303259 (C2C12) and GSE303260 (MSC). The datasets can be accessed via the following GEO link : https://eur03.safelinks.protection.outlook.com/?url=https://3A%2F%2Fwww.ncbi.nlm.nih.gov%2Fgeo%2Fquery%2Facc.cgi%3Facc%3DGSE303260&data=05%7C02%7Cconstance.pirson%40uclouvain.be%7C4dc4199535d54880597908dde9551d08%7C7ab090d4fa2e4ecfbc7c4127b4d582ec%7C1%7C0%7C638923273972140051%7CUnknown%7CTWFpbGZsb3d8eyJFbXB0eU1hcGkiOnRydWUsIlYiOiIwLjAuMDAwMCIsIlAiOiJXaW4zMiIsIkFOIjoiTWFpbCIsIldUIjoyfQ%3D%3D%7C0%7C%7C%7C&sdata=z3EL3o8uH9EcUOSlEn%2F94BBTjWCDfw1ACzDPgGhyu8I%3D&reserved=0 using the secure tokens ebipyiwutvszjut (C2C12) and gfqpyoywxfkddwv (MSC). All other data supporting the findings of this study are available from the corresponding author upon reasonable request.

## Notes

https://www.ncbi.nlm.nih.gov/geo/query/acc.cgi?acc=GSE303259

### Competing Interest Statement

The authors have declared no competing interest.

