## Supplementary Table 1 and Figures 1 to 11 for "*SRF* fusion oncogenes encode constitutively activated chimeric transcription factors in myoid soft tissue tumors"

### SUPPLEMENTARY DATA

#### Supplementary Table 1. Primers used for cloning and site-directed mutagenesis.

| | | |
|---|---|---|
| SRF exon4+stop | fw | TCCAGCGGGACATGAGTGACGCTGCCC |
| | rv | GGGCAGCGTCACTCATGTCCCGCTGGA |
| SRF S162D | fw | GCGCTACACGACCTTCGACAAGAGGAAGACGGGC |
| | rv | GCCCGTCTTCCTCTTGTCGAAGGTCGTGTAGCGC |
| SRF V194E | fw | AGTGAGACAGGCCATGAGTATACCTTTGCCACC |
| | rv | GGTGGCAAAGGTATACTCATGGCCTGTCTCACT |
| FOXO1 ΔTAD | fw | ACGTCATGATGGGCCCTTGAAATTCGGTCATGTCAAC |
| | rv | GTTGACATGACCGAATTTCAAGGGCCCATCATGACGT |
| RELA ΔTAD | fw | GATGATGAAGACCTGTGAGGGGCCTTGCTTGGC |
| | rv | GCCAAGCAAGGCCCCTCACAGGTCTTCATCATC |
| ICA1L Δex13 | fw | TCACAGAGTCCAAAGAAATTAACAAGATGATCCCCCAACAATGG |
| | rv | CCATTGTTGGGGGATCATCTTGTTAATTTCTTTGGACTCTGTGA |
| ICA1L Δex12-13 | fw | GCTGGAGCGTTCAACTGAAACTGGGTCTCCCAA |
| | rv | TTGGGAGACCCAGTTTCAGTTGAACGCTCCAGC |
| ICA1L Δex11-12-13 | fw | GAGCACCTCAGTTTTCTAACTCTGAAAATTGAGTTGCAAAAGATCTAC |
| | rv | GTAGATCTTTTGCAACTCAATTTTCAGAGTTAGAAAACTGAGGTGCTC |
| ICA1L Δex11 | fw | CACCTCAGTTTTCTAACTCTGAAAATAACTGGGTCTCCCA |
| | rv | TGGGAGACCCAGTTATTTTCAGAGTTAGAAAACTGAGGTG |
| ICA1L Δacidic motif | fw | TCTGAAAATGTTGCAAAAGATCTACCTGTACTGAACAACCTCCTAA |
| | rv | TTAGGAGGTTGTTCAGTACAGGTAGATCTTTTGCAACATTTTCAGA |
| SRF ΔMADS | fw | CAAGCCGGGTAAGAAGTCTCCACCCCGTTCA |
| | rv | TGAACGGGGTGGAGACTTCTTACCCGGCTTG |
| SRF ΔMADS β1+2 | fw | CTGTCCACGCTGACAGGGACAACCCGAAAACT |
| | rv | AGTTTTCGGGTTGTCCCTGTCAGCGTGGACAG |
| XbaI-ICA1L ex10-13-XbaI | fw | ATGTTCTAGACAAACAAAGATCACAATGAA |
| | rv | ACATTCTAGATCAAGCATTAAGAAGTTCAT |
| PDGFRβ N666K | fw | CCTGAACGTGGTCAAGCTGTTGGGGGC |
| | rv | GCCCCCAACAGCTTGACCACGTTCAGG |
| PDGFRβ L658P | fw | GATCATGAGTCACCCTGGGCCCCCACCTGA |
| | rv | TCAGGTGGGGGCCCAGGGTGACTCATGATC |
| PDGFRβ JM full | fw | TCCGATGGAAGGTGATTGAGTCTGTGAGCTCTG |
| | rv | TCTCGTAACGTGGCTTAGATGATGCTGTCAGG |
| NotI insertion | fw | GGGAGACCCAAGCTGGCTAGCGTTTAAGCGGCCGCTTGCCACCATGTAC |
| in pcDNA3.1-SRF | rv | GTACATGGTGGCAAGCGGCCGCTTAAACGCTAGCCAGCTTGGGTCTCCC |
| NotI insertion | fw | CGCCGGAATTAGATCTCTCGCGGCCGCCGAATTCTACCGGGTAGGGG |
| in pMSCV-puro | rv | CCCCTACCCGGTAGAATTCGGCGGCCGCGAGAGATCTAATTCCGGCG |

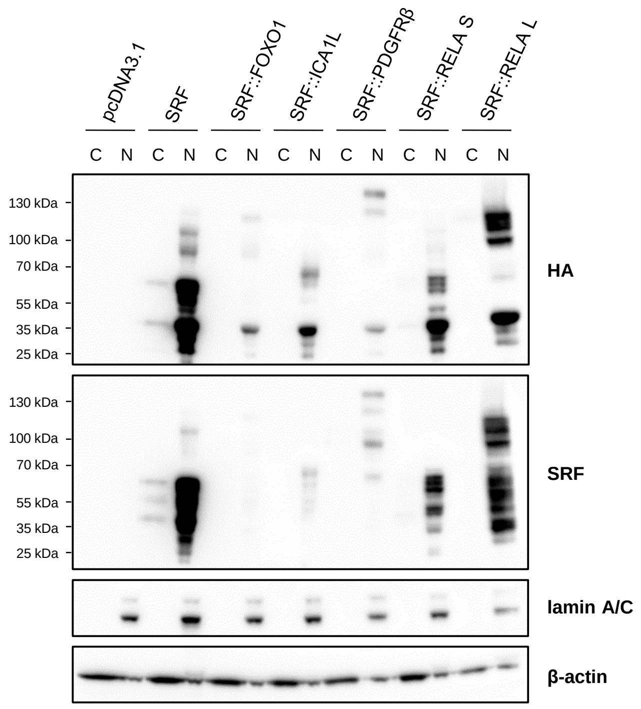


**Supplementary Figure 1. SRF fusions are localized in the cell nucleus.** C2C12 cells were transfected with SRF fusions, and expression was analyzed in cytoplasmic (C) and nuclear (N) extracts by western blot with anti-HA and anti-SRF antibodies (N=1).

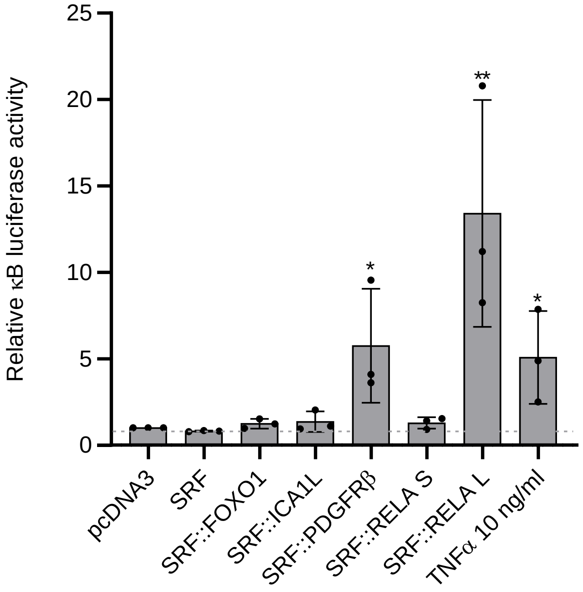


**Supplementary Figure 2. The long version of SRF::RELA activates NF-κB transcriptional activity in an RHD-dependent manner.** C2C12 were co-transfected with SRF fusions, a pκB-luciferase reporter vector and an internal control. After 24h of transfection, luminescence was measured in cell lysates. The mean of three independent experiments is represented with SEM (One-way ANOVA Kruskal-Wallis test with *$p < 0.05$; **$p < 0.01$).

**A**

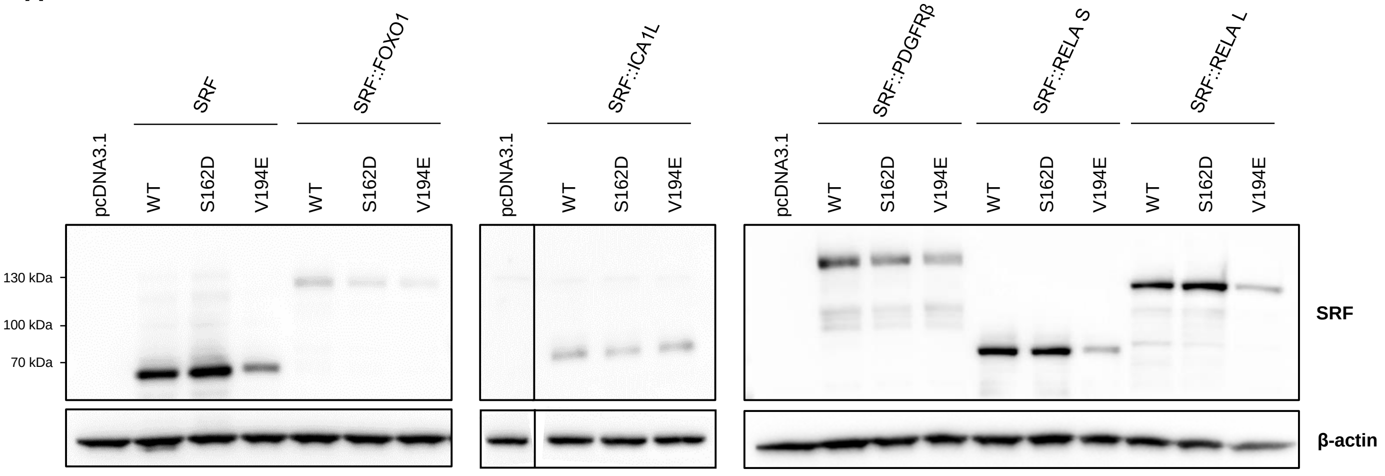


**B**

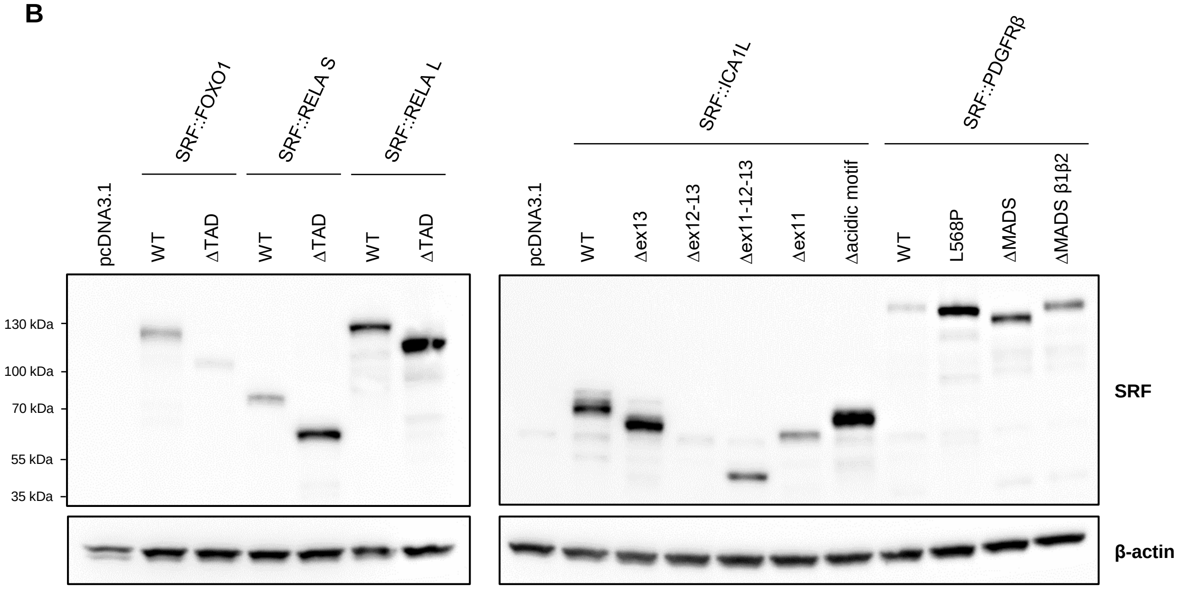


**Supplementary Figure 3. Expression of mutated and truncated SRF fusions used in luciferase assays.** C2C12 cells were transfected with SRF fusions, and expression was analyzed by western blot with anti-SRF antibody (N=2).

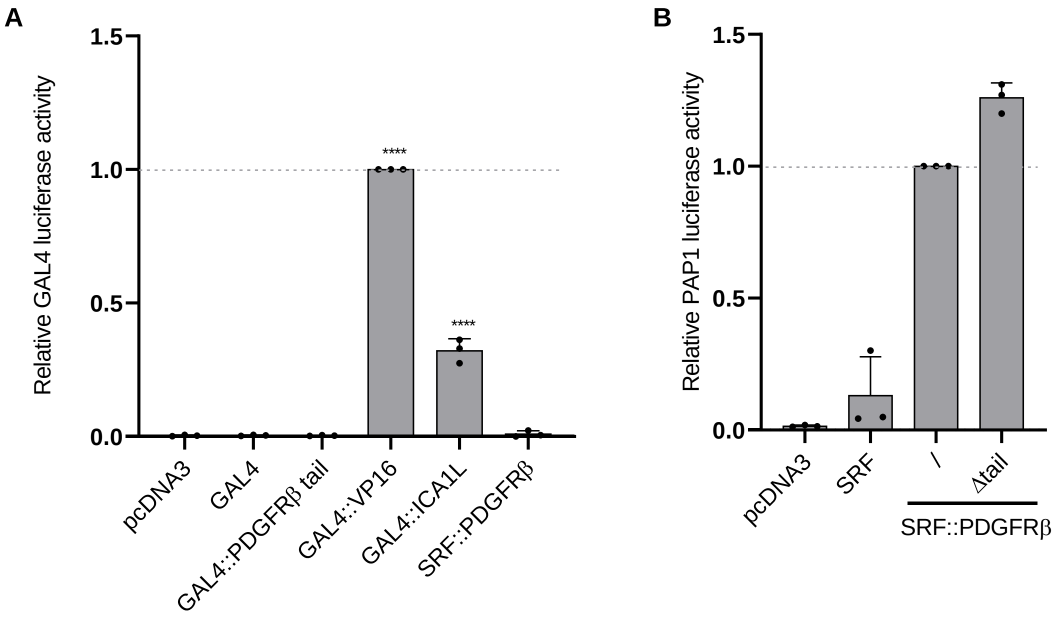


**Supplementary Figure 4. SRF::PDGFRβ fusion is devoid of cryptic transactivation domain.** (A) HEK293T were co-transfected with GAL4 fused with the C-terminus tail of PDGFRβ (the last 119 amino acids) or the C-terminus of ICA1L (exons 10-13), pGL-TK-GAL4-luc reporter vector and an internal control. After 24h of transfection, the luminescence and the β-galactosidase activity were measured in cell lysates (N=3). (B) HEK293T were co-transfected with SRF::PDGFRβ mutants, pPAP1-luc reporter vector and an internal control. After 24h of transfection, luminescence and the β-galactosidase activity were measured in cell lysates (N=3). The mean of three independent experiments is represented with SEM (One-way ANOVA test with ****$p < 0.0001$).

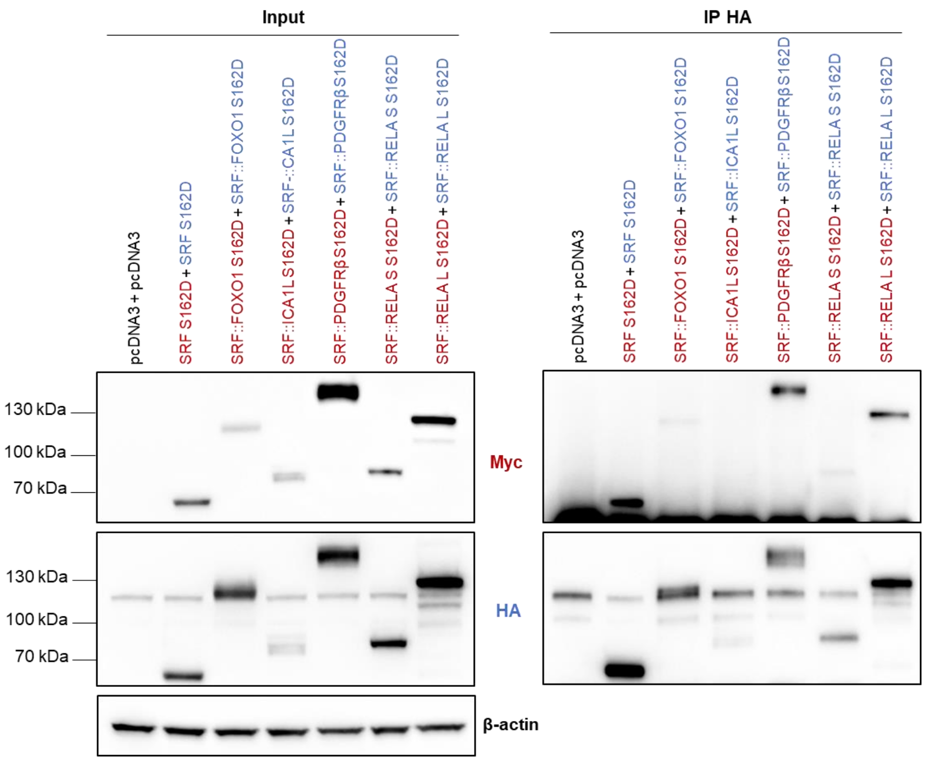


**Supplementary Figure 5. Dimerization of SRF fusions occurs independently of DNA binding.** HEK293T cells were co-transfected with HA- and/or Myc-tagged SRF fusions with a S162D mutation before protein extraction. SRF WT or fusions were immunoprecipitated with anti-HA antibody and analyzed by western blot with anti-HA and anti-Myc antibodies (N=3).

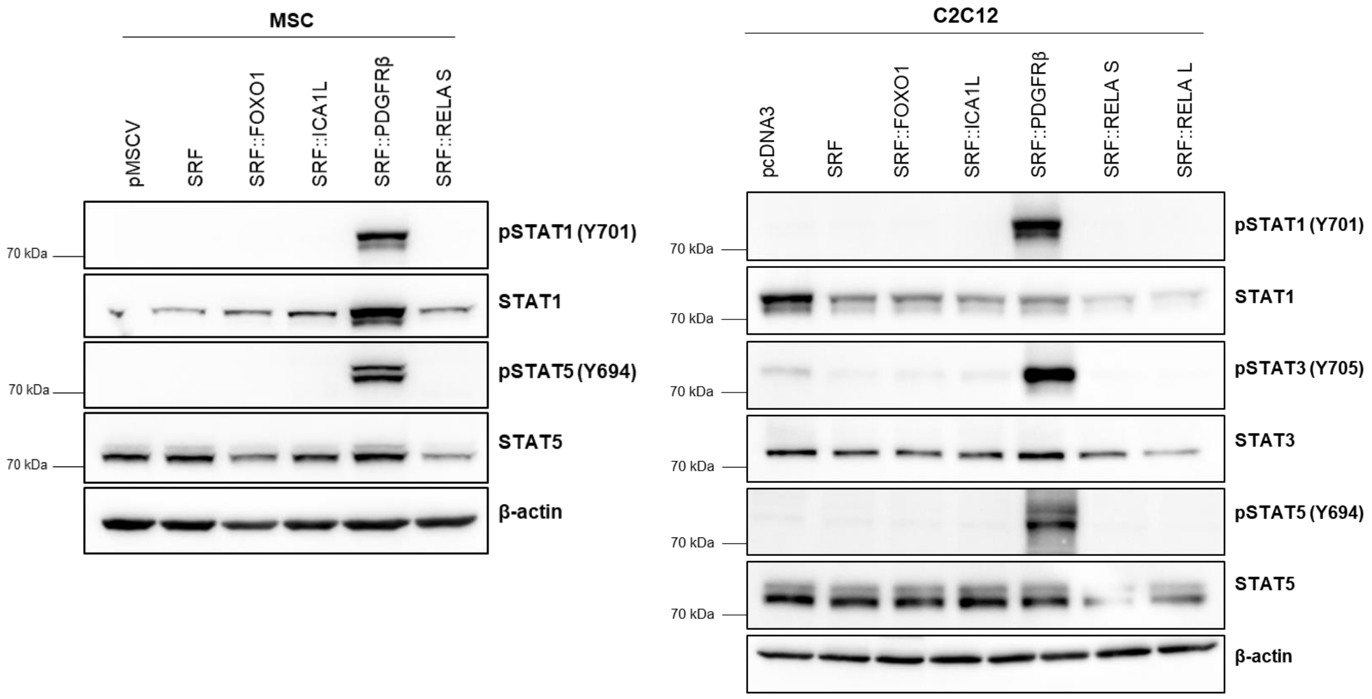


**Supplementary Figure 6. STAT phosphorylation by SRF::PDGFRβ in transduced MSC and transfected C2C12.** Protein expression and STAT phosphorylation after retroviral infection of MSC (N=3) and transfection of C2C12 (N=2) were analyzed by western blot.

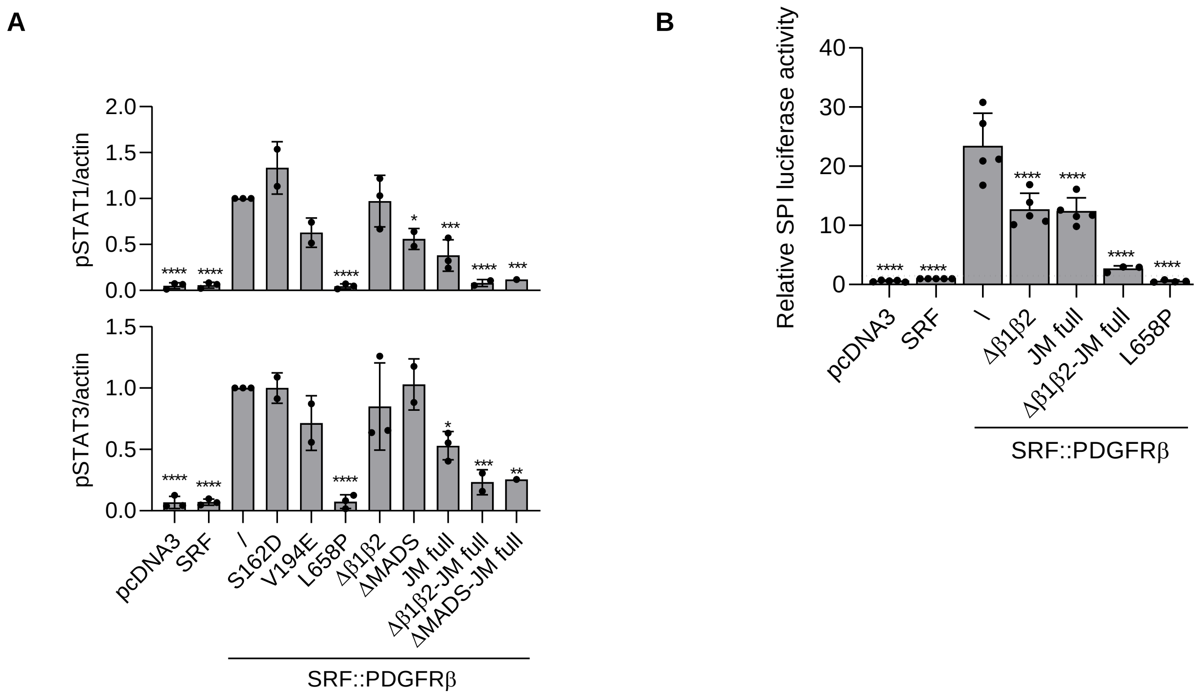


**Supplementary Figure 7. SRF::PDGFRβ activity relies on dimerization and JM domain truncation.** (A) STAT1 and STAT3 phosphorylation was quantified using Bio1D (Vilber) software and normalized to the SRF::PDGFRβ condition. Actin was used instead of STAT1 for normalization because active STAT1 is known to upregulate the STAT1 gene. (B) HEK293T were co-transfected with pcDNA3.1-SRF fusions, pSPI-luciferase reporter vector (containing STAT5-responsive elements) and an internal control. After 24h of transfection, luminescence and β-galactosidase activity were measured in cell lysates (N=3). The mean of three independent experiments is represented with SEM (One-way ANOVA test with *$p < 0.05$; **$p < 0.01$; ***$p < 0.001$; ****$p < 0.0001$).

**MSC**

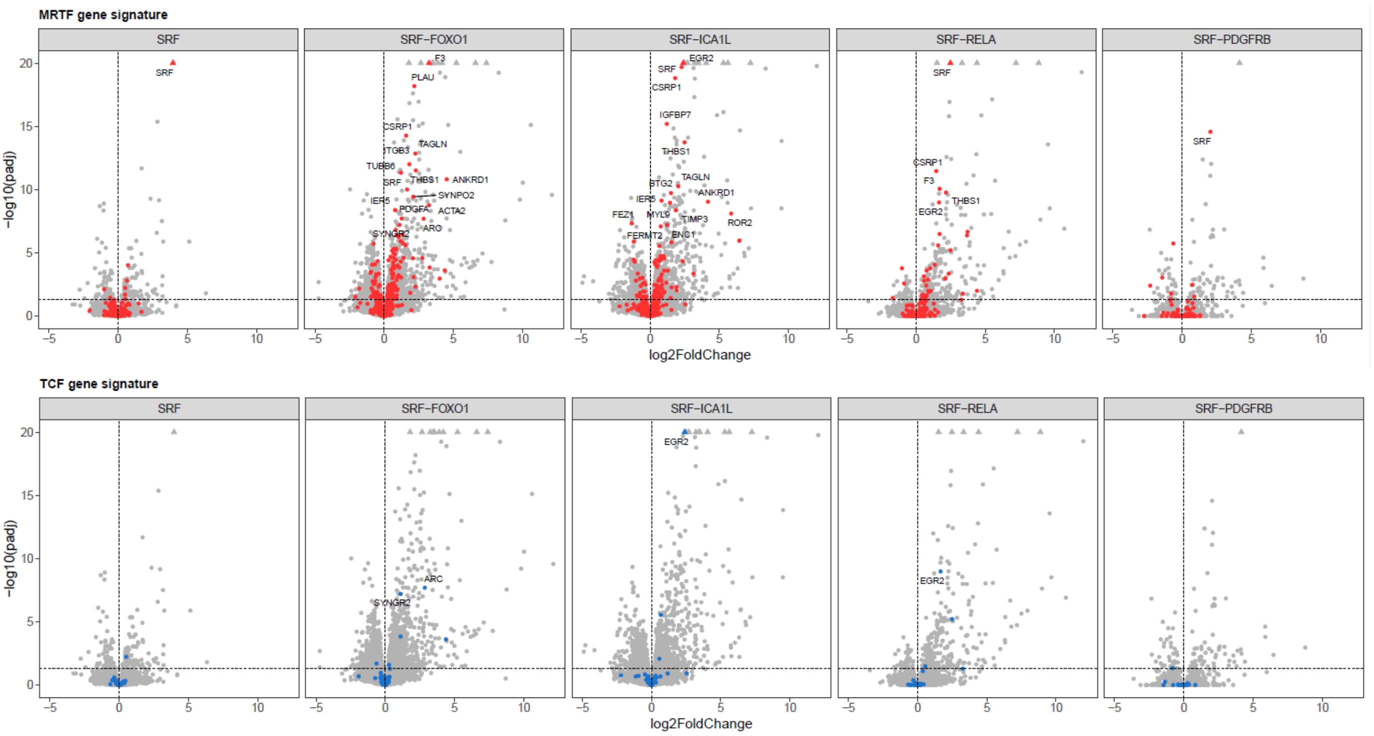


**C2C12**

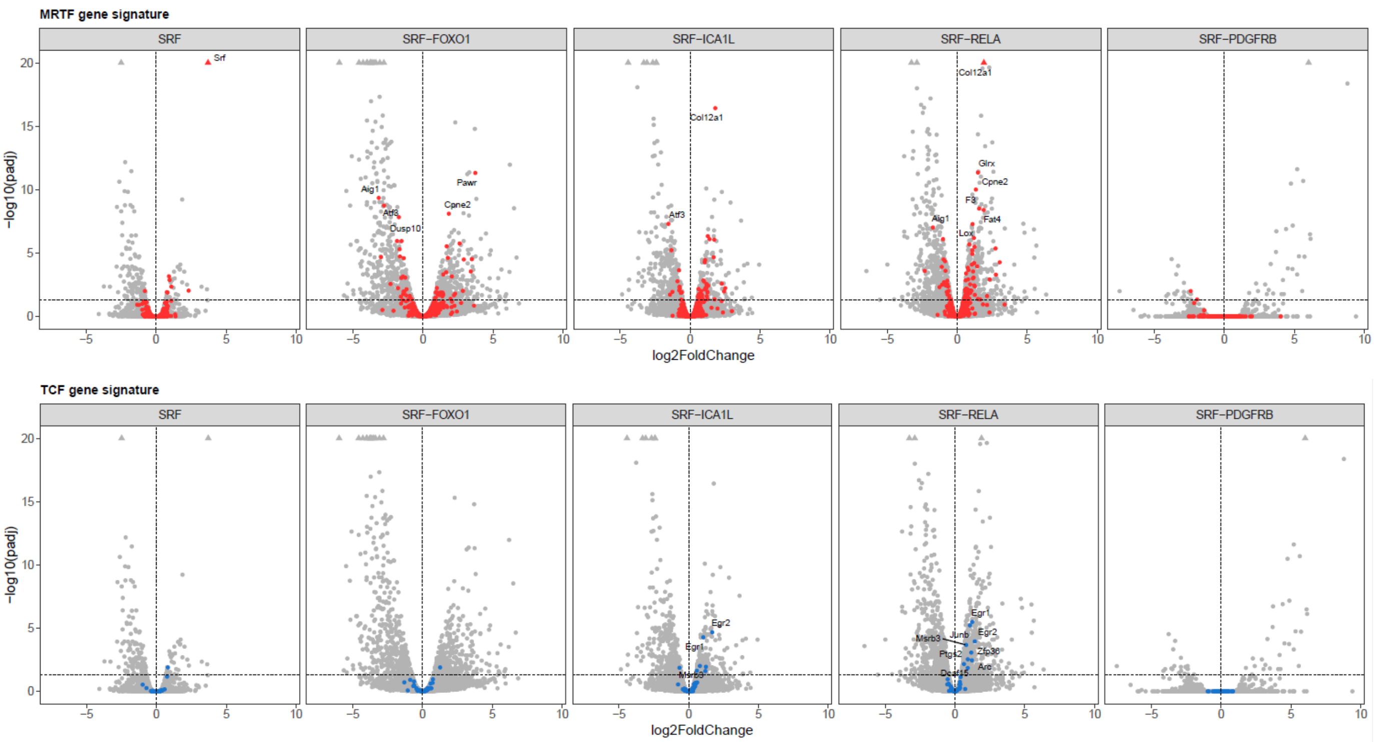


**Supplementary Figure 8. TCF and MRTF target gene expression profiles in MSC and C2C12.** Impact of SRF WT and fusions on SRF/MRTF and SRF/TCF gene expression in MSC and C2C12 cell lines measured by RNA-sequencing. Selected target genes are indicated in red. Fold changes are compared to cells transduced with the empty control vector pMSCV.

**A**

**MUSCLE STRUCTURE DEVELOPMENT**

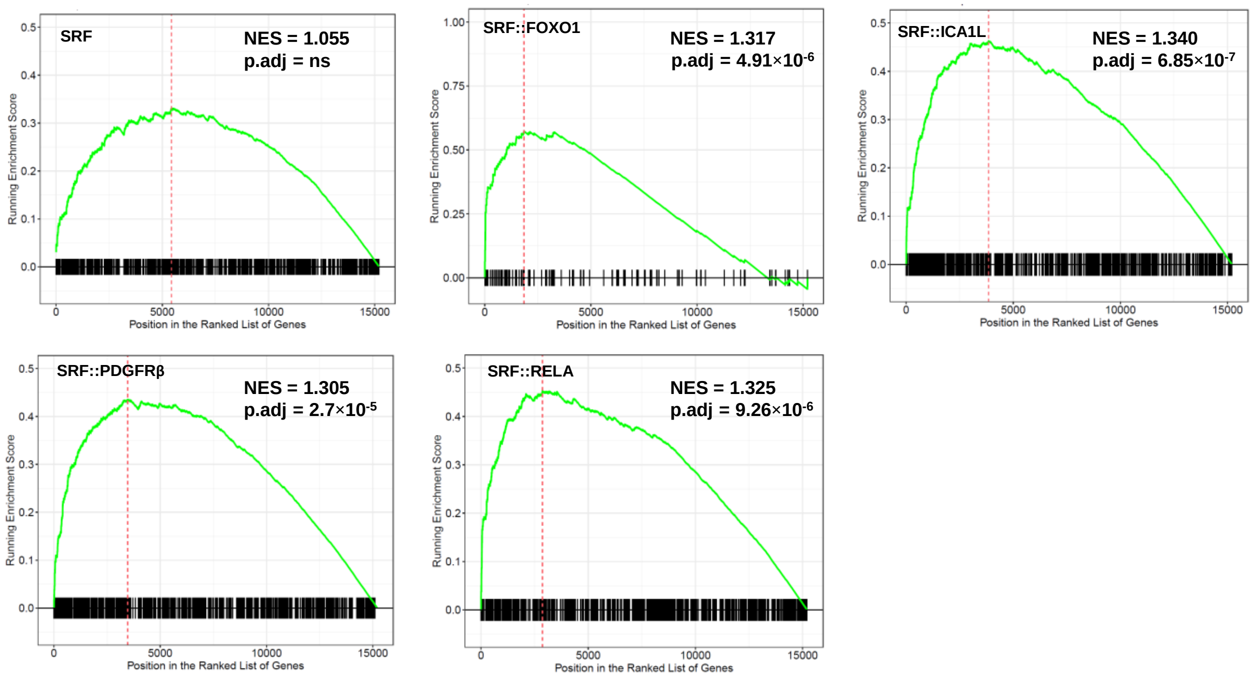


**B**

**Hallmark INTERFERON ALPHA** **Hallmark INTERFERON GAMMA**

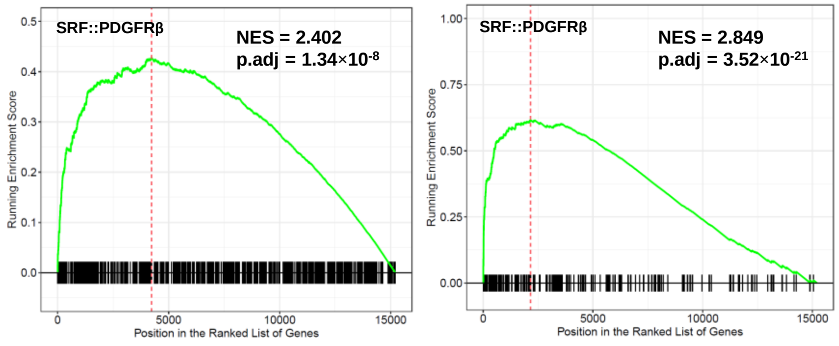


**Supplementary Figure 9. Distinct gene set enrichment signatures in MSC expressing SRF fusions** (A) GSEA analysis revealed that “muscle structure development” gene sets are enriched in SRF fusions-transduced MSC cells. (B) GSEA analysis revealed that « hallmark interferon alpha response » and « hallmark interferon gamma response » gene sets are enriched in SRF::PDGFRβ-transduced MSC cells.

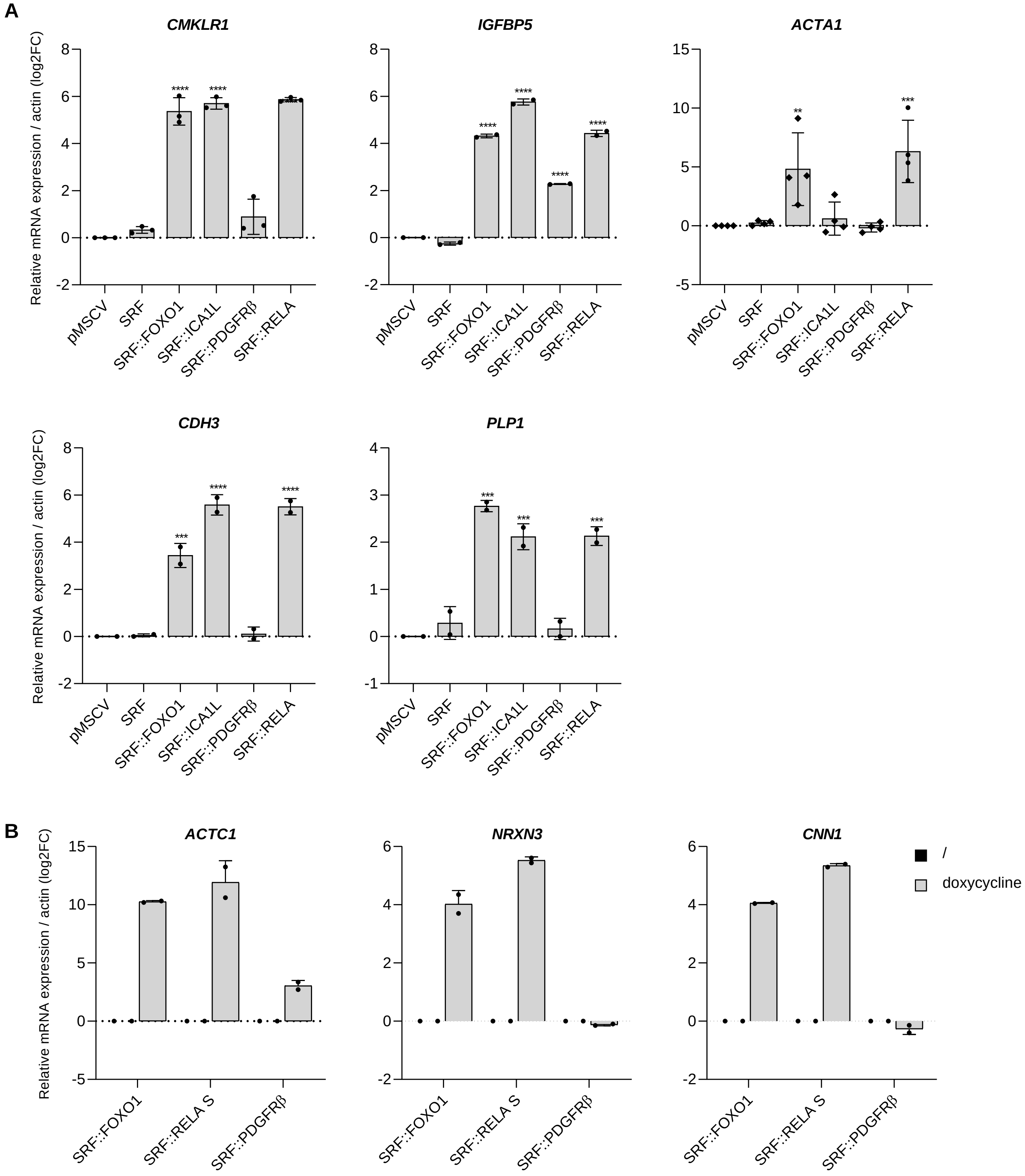


**Supplementary Figure 10. Validation of RNA-seq significantly upregulated genes by RT-qPCR in MSC**. RT-qPCR analysis of genes identified as significantly differentially expressed in the RNA-seq dataset. Relative expression was calculated using the ΔΔCt method and converted to $\log_2$ fold change. The mean of three independent experiments is represented with SEM (One-way ANOVA test with $^{**}p < 0.01$; $^{***}p < 0.001$; $^{****}p < 0.0001$).

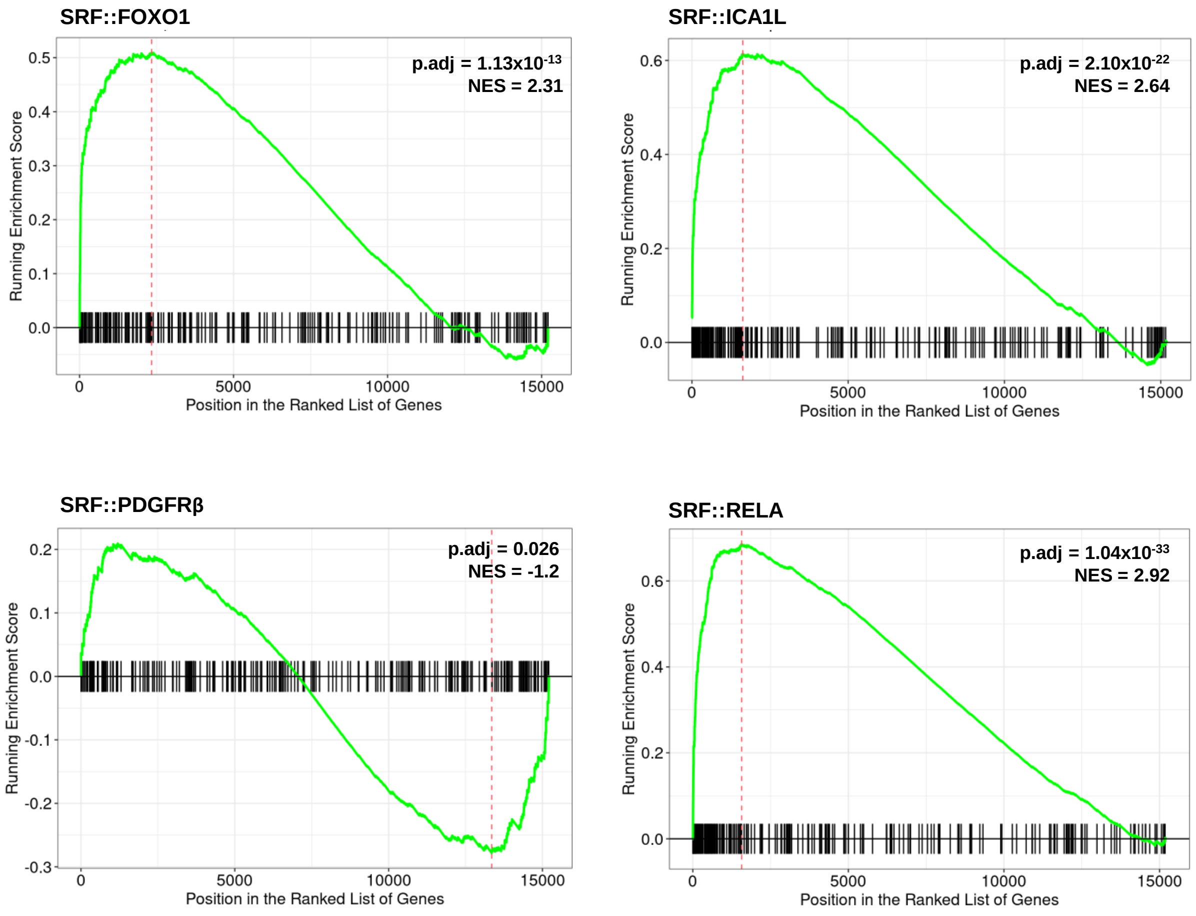


**Supplementary Figure 11. Enrichment plots from Gene Set Enrichment Analysis (GSEA).** GSEA results showing the enrichment of genes upregulated in SRF fusion–positive tumors (n=38), defined by a Bonferroni-adjusted p-value < $1\times10^{-5}$ and a fold change ≥ 3, within the transcriptome of MSC expressing SRF fusions. The green curve represents the running enrichment score, and vertical lines indicate the position of individual genes from the tested gene set within the ranked list.
